# Diversity and single-cell activity of endolithic microbes in sediment-hosted carbonate nodules within and below the sulfate-methane transition zone

**DOI:** 10.1101/2025.02.26.640391

**Authors:** Sergio A. Parra, Magdalena J. Mayr, James Mullahoo, Laura K. Quinn, Rebecca L. Wipfler, Victoria J. Orphan

## Abstract

Authigenic carbonate concretions (‘nodules’) precipitate in marine seep sediments as a result of anaerobic oxidation of methane (AOM). These rocks host active endolithic microbial communities and persist as important methane sinks. Still, how these communities and their activity differ from those in adjacent seep sediments, particularly as a function of proximity to the sulfate-methane transition zone (SMTZ), remains understudied. Here, we sampled sediments and nodules within and below the SMTZ (0-57 cm deep) at four active deep-sea seep areas in Santa Monica Basin, CA. Measurements of high nodule porosities (43-51%) coupled to strong similarities between sediment and nodule 16S rRNA-based community profiles, including below the SMTZ, suggest continued perfusion and exchange between buried nodules and surrounding sediment. Shared, depth-dependent transitions in methanotrophic taxa (ANME-1, ANME-2, ANME-3) and methanogenic taxa (*Methanofastidiosales*) below the SMTZ were also consistent with trends in porewater methane and sulfate concentrations, porewater DIC, and nodule δ^13^C values — underscoring the impact of different geochemical conditions on community structure and suggestive of under-characterized physiological plasticity in ANME-1. Laboratory-based BONCAT incubations of nodules within the SMTZ over ∼14-weeks revealed active sulfide production and translationally active endolithic microorganisms. However, cells from parallel nodule incubations recovered below the SMTZ showed weak-to-negligible BONCAT-based activity despite similar cell abundances — suggestive of low activity on shorter timescales or dormancy. Together, these data challenge the interpretation of passively recorded microbiological signatures in seep sediment-hosted carbonate nodules and expand our understanding of how these endolithic communities may be actively shaped by past and present conditions.

**Importance:** This study advances earlier investigations of microbial communities in buried seep carbonate nodules by integrating microbiological profiles of nodules and sediments, sediment geochemistry, single-cell activity measurements, and nodule mineral, geochemical, and physical characteristics within and below the sulfate-methane transition zone in deep-sea methane seeps. This approach allows us to view how nodule-hosted, endolithic microbial communities change relative to their surrounding sediments across multiple geochemical contexts and better understand how formation history and environmental conditions might affect community identity and metabolic function. Results indicate that the seep nodule and surrounding sediment communities are closely linked across diverse geochemical conditions. This connectivity between sediments and carbonate nodules is distinct from that observed in exhumed seep carbonates, with implications regarding how microbial community composition within these nodules are interpreted, suggesting that instead of a passive recorder of the communities at the time of formation, these nodules appear to retain diverse, metabolically viable communities.

## Introduction

Authigenic carbonates at marine methane seeps form as the result of microbially-mediated anaerobic oxidation of methane (AOM) in sediments, which draws down a substantial amount of the subsurface methane produced at these seeps (1, 2). Because these deposits persist over geologic timescales, these carbonates have been interpreted as a record of past AOM activity in ancient marine environments (3–7). However, several lab-based measurements of elevated methane oxidation rates in recovered seep carbonates demonstrate that these rocks continue to host viable microorganisms and AOM activity, representing a persistent methane sink whose activity and ecology remains under-studied compared to seep sediments (8–11).

In particular, vuggy carbonate concretions, or ‘nodules’ observed in seep sediments (12–15) have not been investigated as thoroughly as denser, seafloor-exposed seep carbonates — whose metabolically diverse, endolithic communities have been observed to differ significantly from sediments and sediment-hosted nodules collected from the same sites, e.g., (16). As such, seep nodules represent a unique opportunity to study seep carbonate microbial community structure and activity in comparison to their host sediments in geochemical context.

For example, AOM in marine sediments primarily occurs within a distinct redox regime known as the sulfate-methane transition zone (SMTZ), where syntrophic activity between anaerobic, methanotrophic (ANME) archaea and sulfate-reducing bacteria (SRB) act to consume porewater methane and sulfate (17–20). This zone of AOM activity varies in depth and thickness largely controlled by methane flux, e.g., (21), transitioning from methane-poor, sulfate-rich overlying sediment horizons to deeper sulfate-poor, methane-rich sediments. The impact of relative proximity to the SMTZ on the endolithic community structure and activity within sediment-hosted carbonate nodules remains poorly characterized, with limited sampling from existing studies focusing exclusively within this zone (14–16, 22). In sediments, this geochemical stratification often results in regimes of distinct ANME/SRB lineages indicative of niche differentiation (22–29), where the observed persistence of certain ANME clades in sulfate-depleted seep sediments remains an area of active investigation (30–32). As such, the SMTZ represents a useful comparative framework in which to study the similarities and differences with co-occurring carbonate nodules across this globally important geochemical transition zone.

Previous work examining nodules recovered from seep sediments reported high similarities in the 16S rRNA diversity and community structure with host sediments (14–16). However, the extent to which these DNA-based community profiles represent extant, metabolically viable communities actively exchanged with host sediment or preserved, ancient sediment communities remains unclear (14, 15), with implications how different traces of AOM are ultimately interpreted and/or preserved in sediment-hosted nodules.

In this study, we expanded the characterization of sediment-hosted authigenic carbonate nodules from four methane seep areas at Santa Monica Mounds 800 and 863 within the Santa Monica Basin (33). Specifically, we integrated the biological and porewater geochemical profiles of 1 m long, ROV-deployed sediment cores within active deep-sea seeps with the structural and microbiological profiles of seep carbonate nodules, effectively capturing changes in both sediment and associated nodules within and well below the SMTZ. Laboratory microcosm experiments were also used to characterize differences in bulk respiration and cellular anabolic activity by the endolithic community across this transition zone using geochemical assays and single cell bioorthogonal non-canonical amino acid tagging (BONCAT) incubations. This combined approach allows us to probe the historical and current capacity of seep carbonate nodules as hosts for continued AOM activity in geochemical context, compared to their surrounding sediments.

## Results

### Sediment and nodule sampling within Santa Monica Basin methane seeps

Nodules and sediment push cores in this study were collected in May 2021 during the WF05-21 Southern California oceanographic expedition on the *R/V Western Flyer* using the *ROV Doc Ricketts*, both owned and operated by the Monterey Bay Aquarium Research Institute (MBARI). 93 samples (7 cores, 82 sediment horizons, 11 nodule horizons) were recovered in total from two active methane seep sites within the Santa Monica Basin: Santa Monica Mound Site 800 (SMM 800: 33.799438N /118.64672E; 805m depth; also known as the NE mound) and Santa Monica Mound Site 863 (SMM 863: 33.7888N /118.6683W; 863 m; also known as the SW mound); (33, 34). At these sites, we investigated 4 distinct seep areas: SMM 800-I, SMM 800-II, SMM 800-III, and SMM 863 (Fig. 1). Within each seep area, 1-2 sediment cores were collected, including 3, 1.22 m long cores (LC) and 4, standard, 30 cm long push cores (PC), with quasi-duplicate (‘paired’) cores taken in close spatial proximity (∼30-50 cm apart) to capture comparable environments, where possible. One sediment core (PC64, SMM 800-II) was collected during a previous cruise and sampled from the same microbial mat within 5 m from the 2021 sampling location (Supplemental Material).

**Fig. 1.**
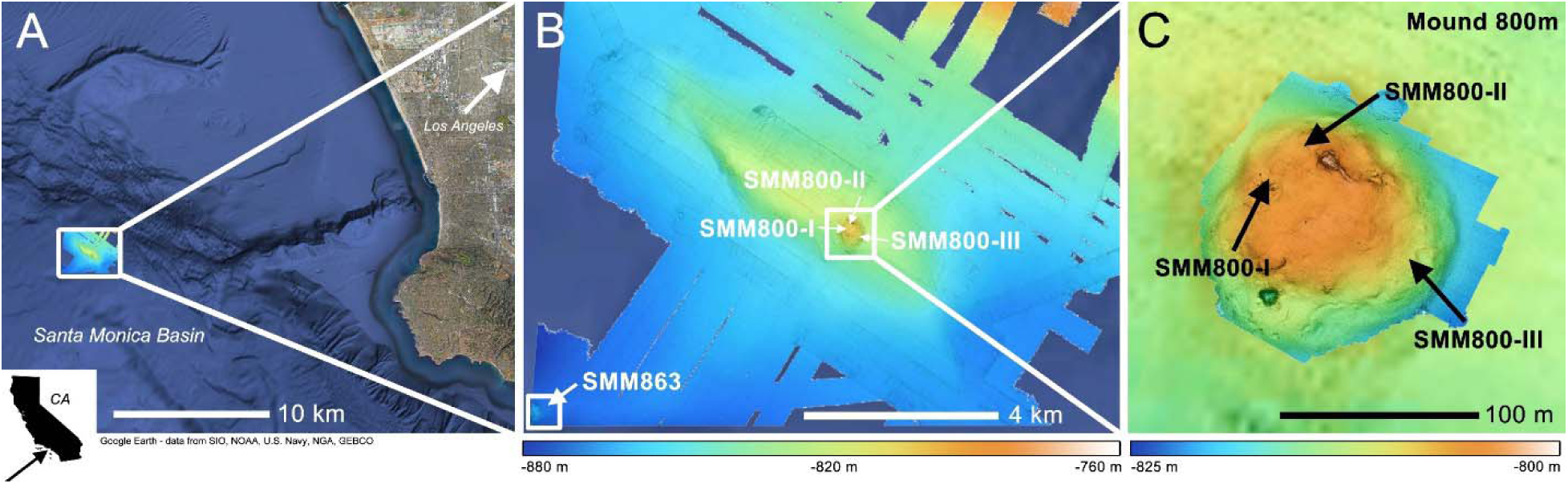
A) General location of the Santa Monica Basin seep sites (NE Mound, 800m, and SW mound, 863m) sampled for this study; B) Multi-beam bathymetry map of the sampling seep areas at SMM 800 and SMM 863; C) higher-resolution bathymetry map of the seep areas sampled at SMM 800: SMM 800-I, SMM 800-II, and SMM 800-III. Bathymetry colors in B) and C) correspond to mapped depth. Bathymetry maps were collected by MBARI in 2018.

### Morphological diversity in seep sediment-hosted nodules

Across both short and long sediment push cores from all four seep areas, we observed distinct depth intervals of hardened, grey-brown calcitic material ranging in size from <CM rubble to >cm concretions, or nodules. Representative nodules from the four seep areas are shown in Fig. 2, with a complete list of the nodule-bearing sediment horizons included in this study summarized in Table S1. We observed strong morphological differences between the nodules recovered from cores at SMM 800 and SMM 863. Nodules from SMM 800 (Fig. 2A-C) were rougher, more irregularly shaped, and featured a greater number of mm to cm-sized voids. These nodules were also less friable, with lobate, grey-brown, coarse-grained growths often covering black, thin bivalve shell fragments. These features were consistent between the three seep areas sampled at SMM 800, as well as down core (7 nodule-bearing horizons total). In contrast, nodules from SMM 863 (Fig. 2D) were less irregularly shaped, with a less porous, finer grained, brown-colored material that resembled packed sediment hosting smaller lithics within. This morphology was also consistently observed between cores and across the five nodule-bearing horizons sampled at SMM 863.

**Fig. 2.**
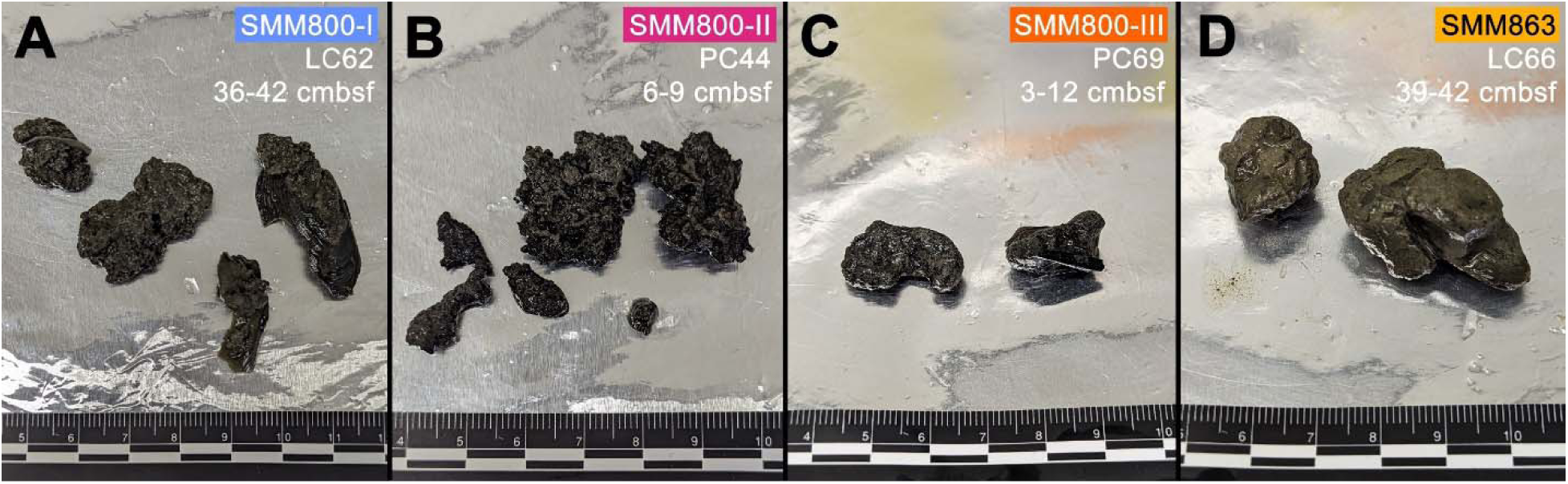
Characteristic nodules recovered from each of the four seep areas in this study; Nodule morphology is remarkably consistent with depth (expressed as centimeters below seafloor, cmbsf); ruler markings are in cm; A) Nodules recovered from LC62’s 36-42-cmbsf horizons at SMM 800-I; B) Nodules from PC44’s 6-9-cmbsf horizon at SMM 800-II; C) Nodules recovered from PC69 (3-12-cmbsf) at SMM 800-III; D) Nodules recovered from the 39-42-cmbsf horizon of LC66 at SMM 863.

### Nodule porosity differences and pore size distributions

Open porosity is defined as the volume of connected pores relative to the volume of bulk solid, effectively representing the pore space that can freely exchange with the surrounding sediment environment. Water intrusion analyses of a subset of nodules (Table 1) revealed that the open porosities of two nodules sampled from SMM 800 (800-I, 800-II) were substantially higher than that of the nodule from SMM 863.

**Table 1.** Water intrusion-derived open porosities for select nodules.

| Seep Area | SMM 800-I | SMM 800-II | SMM 863 |
| --- | --- | --- | --- |
| Core ID | LC62 | PC44 | LC66 |
| Core Horizon (cmbsf) | 36-42 | 9-12 | 39-42 |
| Open Porosity (%) | 51.36±0.84 | 49.52±1.33 | 43.03±0.78 |

Hg-porosimetry enabled the measurement of pore size distributions between 6 and 350 μm, revealing several pore-size regimes within a SMM 863 nodule collected at depth (39-42 cmbsf), including peaks corresponding to 11, 30, and 300 μm, and a slightly higher proportion of pores with sizes >100 μm (Fig. 3). A similarly deep-sourced nodule from SMM 800-I had fewer, although more evenly distributed pore sizes in the observed range. We also saw a smaller peak of pore diameters at 90 μm and fewer pore diameters above 100 μm compared to the nodule from SMM 863. A shallower nodule from SMM 800-II (9-12 cmbsf) also had a relatively low, but even pore-size distribution below 100 μm, in addition to the occurrence of pore sizes between 100 and 250 μm, with minor peaks at 300 and 350 μm.

**Fig. 3.**
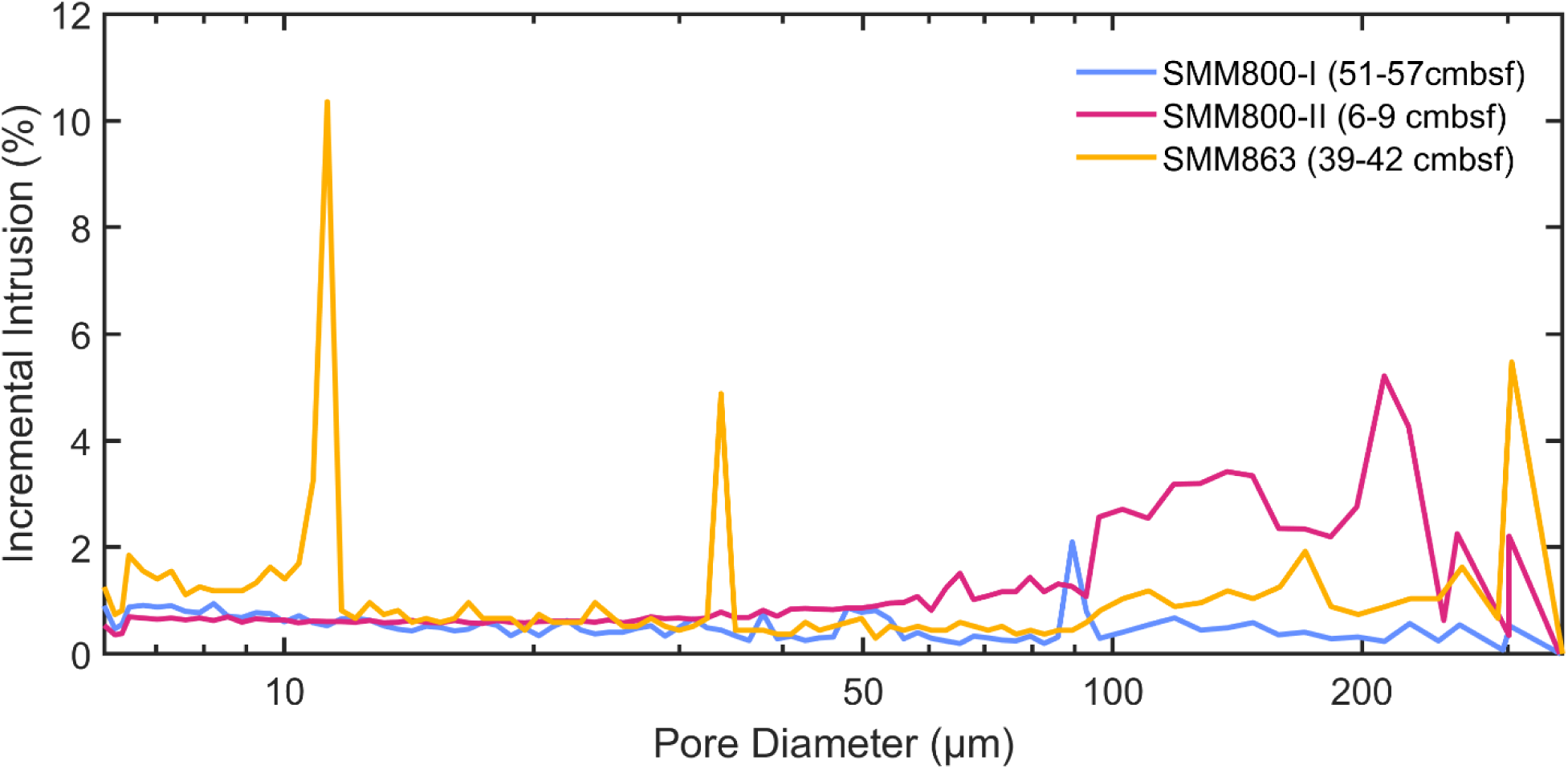
Hg-porosimetry curves detailing pore diameter versus incremental intrusion for three nodules sampled from SMM 800-I, SMM 800-II, and SMM 863. Pore sizes observable with porosimetry range from 6- to 350-μm. The three nodules shed information on the three distinct nodule-featuring regions identified in this data set: 1) SMM 800 deep sediment (LC62), 2) SMM 800 shallow sediment (PC44), and 3) SMM 863 deep sediment (LC66)

### Aragonite dominates nodule carbonate mineralogy, with increased Mg-calcite fractions in deeper nodules

Bulk powder XRD-derived mineralogies showed SMM 800 nodules were largely composed of aragonite (>78.2%), with minor contributions from Mg-calcite, calcite, and dolomite (Table 2). Conversely, nodules from SMM 863 were more aragonite-poor (<43.1%) and featured greater contributions from Mg-calcite (45.1%, 65.1%) than the nodules from SMM 800. Collectively, however, we did observe a greater proportion of Mg-calcite in the nodules sourced from deeper sediment horizons.

**Table 2.** Key mineral phases for select nodules, as determined by XRD.

| Seep Area | Core ID | Depth (cmbsf) | Aragonite | Mg-Calcite<sup>a</sup> | Calcite | Dolomite | Quartz |
| --- | --- | --- | --- | --- | --- | --- | --- |
| **SMM 800-I** | LC62 | 24-27<br>48-51 | 85.2%<br>78.2% | 4.5%<br>11.3% | 0.6%<br>3.8% | 3.3%<br>0.03% | 6.4%<br>6.7% |
| **SMM 800-II** | PC44 | 9-12 | 90.2% | 2.1% | 2% | 1.6% | 3.3% |
| **SMM 800-III** | PC69 | 3-12 | 97% | 0.1% | 1.3% | 0.8% | 0.9% |
| **SMM 863** | LC74 | 33-39 | 43.1% | 45.1% | 0.5% | 4.8% | 0.5% |
|  | LC66 | 39-42 | 26.1% | 65.1% | 7.1% | 1.6% | 0.03% |

a: Mg-Calcite corresponds to $(Ca_{0.9}Mg_{0.1})O_3$ , as measured by (35)

### Geochemical localization of the SMTZ and **δ**^13^C trends in porewater DIC and nodules

To characterize the geochemical environments at SMM 800 and SMM 863, we focused on the vertical trends of three characteristic porewater species that inform the sulfate-methane transition zone (SMTZ): methane (CH_4_), sulfate (SO ^2-^), and sulfide (H_2_S), shown in Fig. 4. At SMM 800-I (LC62) collected within a sulfur-oxidizing microbial mat (Fig. S1), porewater extracts revealed high AOM activity, with a near-complete drop in sulfate concentration by 6-9 cmbsf, which remained low down core (Fig. 4A). This drop was matched by a concurrent rise in porewater sulfide and methane, although we observed more dynamic behavior in the sulfide and methane concentrations with depth, including a notable decrease in both species below their peaks at 6-9 cmbsf. SMM 800-II (PC64) showed a similar sulfate profile, reaching its minimum by 3-4 cmbsf, with values measured at ∼3 mM to the base of the short core at 9 cmbsf (Fig. 4B). Similarly, this drop was accompanied by a steep rise in porewater methane and sulfide, although as with SMM 800-I, we observed a subsequent drop in concentration below the 3-4 cmbsf horizon. At SMM 800-III (PC43), also collected in a sulfur-oxidizing microbial mat (Fig. S3), the peak in AOM activity also occurred at shallower depths (4-5 cmbsf), and, distinct from SMM 800-II, sulfate concentration dropped below detection below this depth (Fig. 4C). At SMM 863 (LC66), only porewater sulfate concentrations were measured. However, compared to the seep areas at SMM 800, we observed a much deeper penetration of sulfate into the sediment, dropping below detection by 21-24 cmbsf in this microbial mat habitat, suggestive of a lower methane flux (Fig. 4D).

**Fig. 4.**
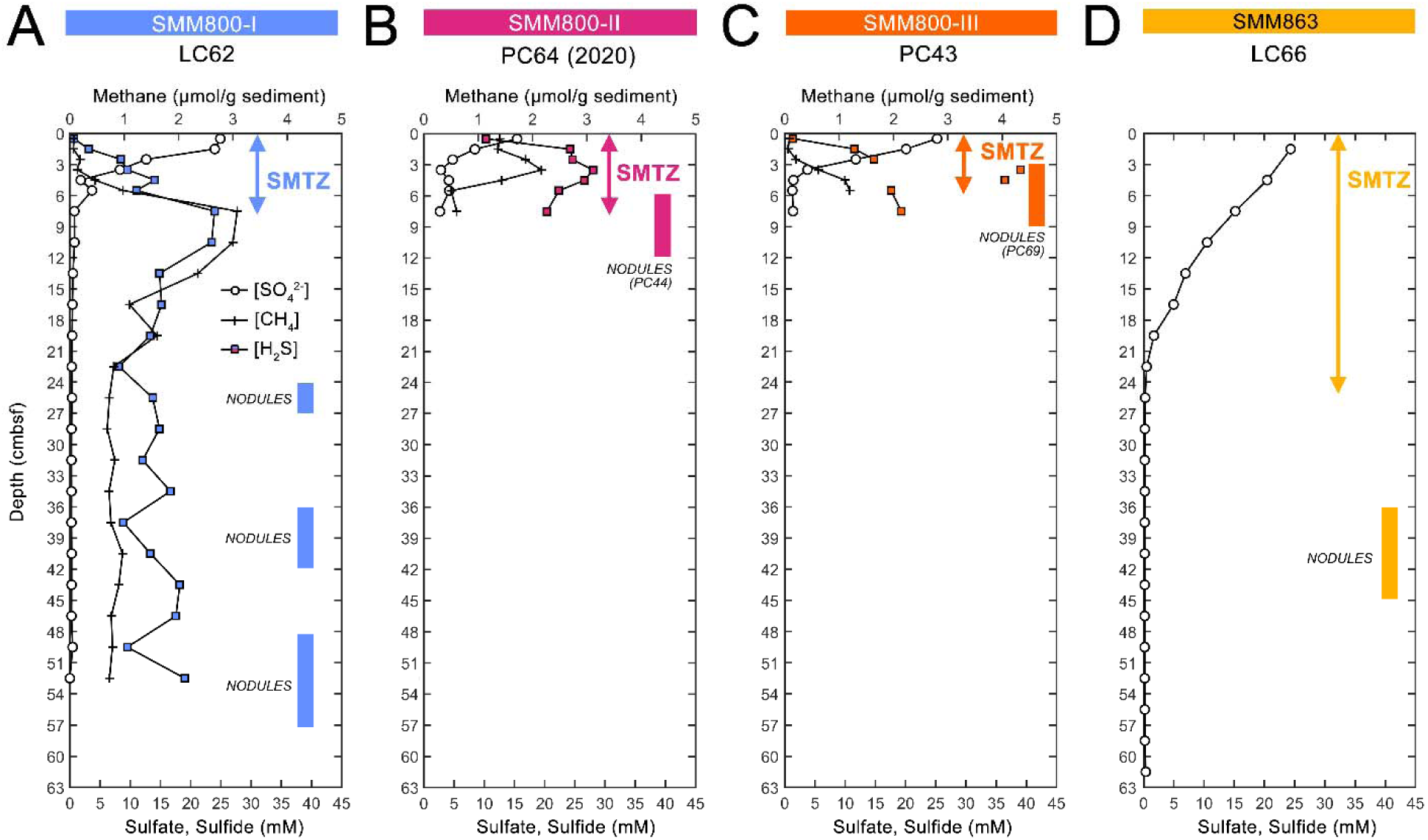
Sulfate, methane, and sulfide concentration profiles collected for sediment porewaters at the seep areas (A-D) investigated in this study. We plot the median depth in centimeters below seafloor (cmbsf), where concentrations represent 3-cm depth horizons. The sulfate-methane transition zones (SMTZ), as determined by near-complete(<<1mM) sulfate drawdown in each core, are also demonstrated with colored arrows. Colored bars represent the horizons where nodules were recovered for further analysis. Nodules recovered from a different, co-located core at the seep area are also specified. PC64 from SMM 800-II was collected a year previously. We note that there are greater uncertainties with measured sulfide concentrations above the detection limit (>25mM, e.g., in PC43 from SMM 800-III). Only sulfate porewater concentration was collected for the core at SMM 863.

Porewater concentrations and the δ^13^C values of dissolved inorganic carbon (DIC, δ^13^C_pw_) in addition to the inorganic carbon isotopic concentration of the nodules (δ^13^C_nod_) were also measured across the four areas in this study (Fig. 5). These data provide additional information into active AOM horizons where highly ^13^C-depleted bicarbonate is produced during methane oxidation and subsequently serves as the source of precipitation of authigenic carbonates (36). At SMM 800-I (LC62), porewater DIC concentrations steeply increased from <10mM to as high as ∼42mM in the shallow subsurface (0-6 cmbsf) before stabilizing down core. Interestingly, thi occurred in two distinct stages at SMM 800-I, which experienced a second rise in DIC to ∼36mM at 15-18 cmbsf to the base of the core at 54 cmbsf (Fig. 5A). At both SMM 800-I and SMM 800-III (PC43), this shallow spike in DIC concentration was paired with the peak in negative δ^13^C_pw_, reaching values as low as -59.20‰, followed by a steady increase in δ^13^C value down core. This indicated a peak in AOM in the shallow sediment horizons, consistent with the depth of the sulfate minima. SMM 800-II (PC64) exhibited a different trend, with comparatively less δ^13^C_pw_ (-46.24‰, Fig. 5B). Notably, both DIC concentration and δ^13^C_pw_ increased with depth, perhaps reflective of a mixture of both heterotrophic sulfate reduction and AOM at this site (37, 38). In comparison, δ^13^C_nod_ exhibited a wide range of values across all four seep areas (-30.09‰ to -53.79‰). Notably, nodules from shallower depths at both SMM 800-I and SMM863 had greater ^13^C enrichment than the nodules recovered deeper in the cores, where the δ^13^C_nod_ at SMM 800-I was slightly more ^13^C-depleted (-46.03‰) than its surrounding porewater (-41.26‰).

**Fig. 5.**
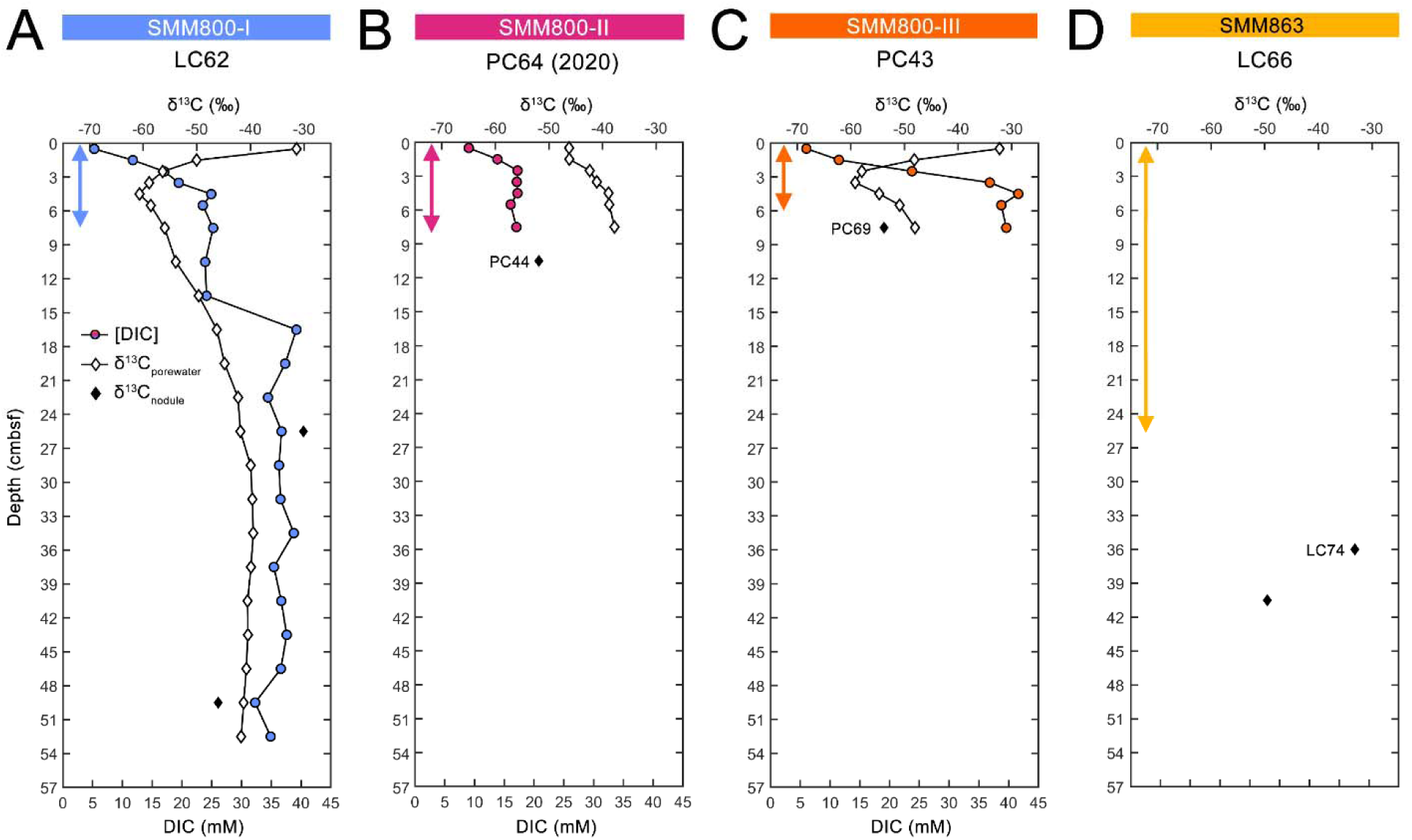
Sediment porewater DIC, δ^13^C_porewater_, and δ^13^C_nodule_ profiles collected at the seep areas investigated in this study. We plot the median depth in centimeters below seafloor (cmbsf), where values represent multiple depth horizons. Because of the <3‰ spread between δ^13^C_nodule_ replicates, values are simply averaged for comparison (individual values are included in Table S2). Colored arrows indicate an estimate for the location of the SMTZ, as in Fig. 4. PC64 from SMM 800-II was collected a year previously. Only δ^13^C_nodule_ was able to be collected for the cores at SMM 863.

### 16S rRNA gene comparison of phylum-level diversity in nodules and host sediments

Dominant microbial phyla across all sampled sediment and nodules (Fig. S6-S12) broadly resembled reports from other carbonate-bearing methane seep environments from Hydrate Ridge, OR and Eel River Basin, CA, which were often dominated by members of the *Halobacteriota* (putative methane-oxidizers largely affiliated with ANME archaea lineages), *Euryarchaeota*, and *Desulfobacterota* phyla (14, 16, 22). Additionally, members of the archaeal *Thermoplasmatota*, *Asgardarchaeota*, and *Crenarchaeota* were consistently recovered above 1% relative abundance. Of the bacterial phyla, *Desulfobacterota* were consistently present, with additional representation by members of the *Chloroflexi* and *Caldatribacteriota* (the latter exclusively associated with JS1, Table S11), and occasional detection (>1%) of taxa belonging to the *Planctomycetota*, *Acidobacteriota*, and *Fermentibacterota*. Notably, members from the *Fermentibacterota,* first characterized in anaerobic digesters by (39), were only substantially recovered in three nodules (1.05-2.28% relative abundance) at both the SMM 800-II and SMM 800-III seep areas and rarely recovered (<<1%) within the associated host sediments (Fig. S9, S11).

### Similar 16S rRNA community profiles between sediments and nodules from shallow sediment horizons

In the shorter push cores from seep areas SMM 800-II and 800-III, *Halobacteriota* lineages consistently present in seep sediments were ANME-1a, ANME-2b, and ANME-2c (Fig. 6). Interestingly, these lineages persisted past the shallow zone of sulfate depletion in these cores. ANME-2c relative abundances were largely consistent with depth through 9-12 cmbsf at both SMM 800-II and 800-III. However, we observed contrasting patterns in ANME-2b and ANME-1a abundances. Deeper horizons in SMM 800-II (PC44) showed a drop in the relative abundance of ANME-2b, concurrent with an increase in ANME-1a below 6 cmbsf. In the cores from SMM 800-III (PC43 and PC69) however, ANME-2b relative abundance and comparatively less ANME-1a were recovered from the deeper horizons, although this trend was much weaker in PC69 (Fig 6C). Within the three nodules recovered from these push cores, the dominant *Halobacteriota* lineages largely resembled those of their parent sediment horizons, where notable exceptions include an enrichment of ANME-1a (>20%) in the nodules from PC44 at SMM 800-II. Still, Shannon-Wiener and Inverse Simpson indices based on archaeal 16S ASV proportional abundances (Table S3) showed that archaeal diversity was greater in the sediment communities from SMM 800-II and SMM 800-III than in the nodules.

**Fig. 6.**
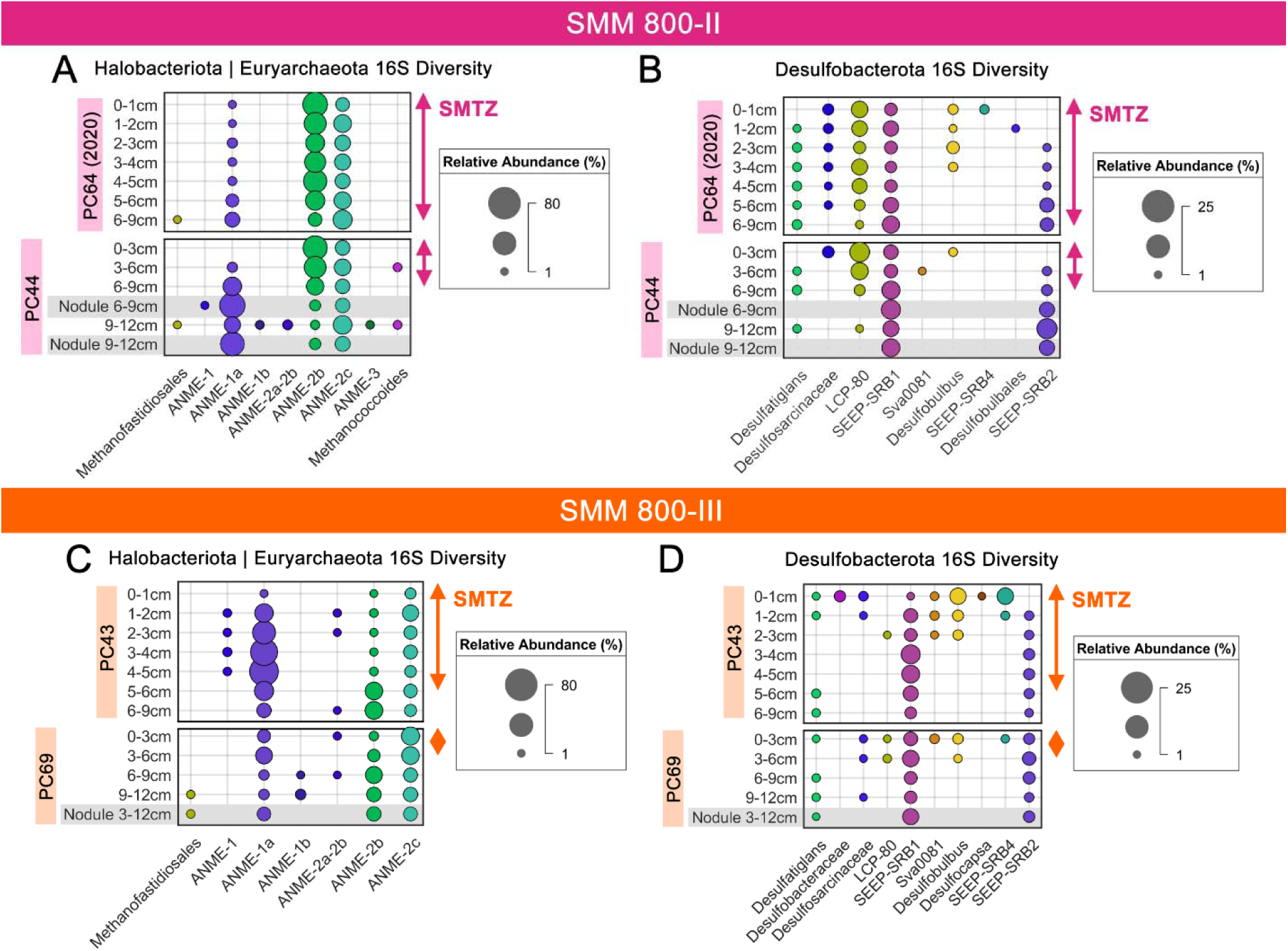
Bubble plots summarizing the depth-resolved, 16S relative abundances for groups within *Halobacteriota* (featuring ANME), *Euryarchaeota (*featuring methanogens, e.g., *Methanofastidiosales*), and *Desulfobacterota* (featuring sulfate-reducing partners to ANME) in PC64 and PC44 at SMM 800-II (A, B), as well as PC43 and PC69 at SMM 800-III (C, D). Where possible, ASVs within these phyla were grouped at the genus-level, or at the next highest taxonomic level possible, and then cut off below a minimum 1% relative abundance. Communities extracted from nodules are highlighted in grey and often represent larger depth horizons. The location of the sulfate-methane transition zone (SMTZ) according to porewater sulfate concentrations from the same or co-located sediment core is indicated with the colored arrows.

Dominant *Desulfobacterota* lineages within these shallow sediment horizons at SMM 800-II and 800-III included two main sulfate-reducing bacterial lineages, Seep-SRB1 and Seep-SRB2, that have been previously documented to form syntrophic associations with ANME (40–43), where their respective abundances appeared to covary with those of their ANME partners (Fig. 6BD). In the sulfate-rich, near-seafloor sediments (0-3 cmbsf) at both seep areas, a large diversity of other putative sulfate-reducing bacterial groups was also detected above 1%. These SRB lineage were largely affiliated with groups not known to directly participate in syntrophic AOM. By comparison, *Desulfatiglans* (not currently known to be involved in AOM) was recovered from both seep areas at low abundances as deep as 12 cmbsf, including within the nodule from PC69 at SMM 800-III. While uncommon in seep ecosystems, this group has been observed in cold seep sediments in the Okinawa Trough that also harbor ANME-1 (29). Shannon-Wiener and Inverse Simpson indices based on bacterial 16S ASV proportional abundances (Table S3) indicated that the sediments surrounding the nodules from these seep areas had higher diversity, as found with the Archaea.

### Deeper sediment and nodule 16S rRNA surveys show persistence of *Halobacteriota* and *Desulfobacterota* and putative methanogenic lineages

Similarly to SMM 800-II and 800-III, SMM 800-I (LC62) had a shallow SMTZ (sulfate depleted by 5-6 cmbsf) and both sediments and nodules were enriched in *Halobacteriota*, *Euryarchaeota*, and *Desulfobacterota* lineages within and well below the sulfate depletion depth. Archaeal diversity was largely dominated by ANME-1a (up to 66.5% relative abundance) which was recovered throughout the core at depths below 3 cmbsf. In the deeper horizons, this group was accompanied by ANME-1b at lower relative abundance (Fig. 7). Notably, within this long core, we also documented two distinct transitions in both archaeal and bacterial community composition, which occurred between 4-27 cmbsf and between 33-57 cmbsf, representing the base of the core.

**Fig. 7.**
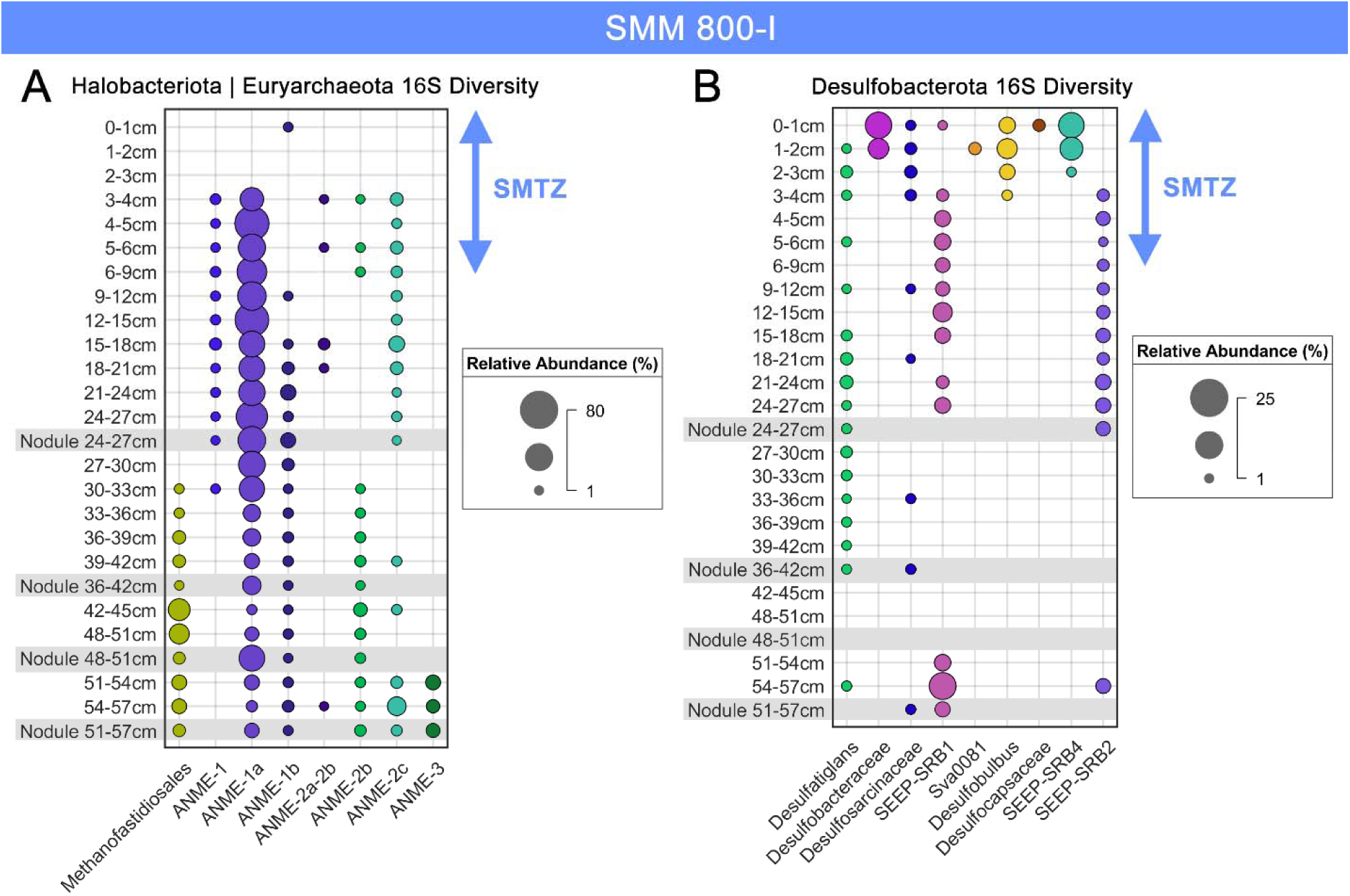
Bubble plots summarizing the depth-resolved, 16S relative abundances for groups within *Halobacteriota*, *Euryarchaeota*, and *Desulfobacterota* in LC62 at SMM 800-I. Where possible, ASVs within these phyla were grouped at the genus-level, or at the next highest taxonomic level possible, and then cut off below a minimum 1% relative abundance. Communities extracted from nodules are highlighted in grey and often represent larger dept horizons. The location of the sulfate-methane transition zone (SMTZ) according to porewater sulfate concentrations from the same or co-located sediment core is indicated with the colored arrows. A) Relative abundances within prominent phyla *Halobacteriota* (featuring ANME) and *Euryarchaeota* (featuring methanogens such as *Methanofastidiosales*). B) Relative abundances within *Desulfobacterota* – the phylum with sulfate-reducing partners to ANME.

The first transition showed a shift from the consistent presence of ANME-2c (1-10% relative abundance), an unclassified ANME-1 lineage, and the putative syntrophic sulfate-reducing bacterial lineages Seep-SRB1 and Seep-SRB2 in the shallow sediment down to 27 cmbsf to an underlying zone ∼24 cm thick where each of these groups, with the exception of sporadic occurrence of ANME-2c, were below detection (Fig. 7). The second transition, occurring between 33-57 cmbsf (well below the SMTZ), was marked by the emergence of ANME-2b and a putative methylotrophic methanogen lineage *Methanofastidiosales* (formerly the WSA2 group, <20% relative abundance) largely dominated by a singular ASV (Table S11). The *Methanofastidiosales* are a poorly characterized group originally described from anaerobic sludge digesters (44) and observed in hydrothermal systems, gas hydrates, methane seeps, and methane-rich coastal sediments (29, 32, 45–48). This ASV was also detected above 1% abundance in the shallower push core sediments from SMM 800-II (PC464, PC44) and 800-III (PC69), including a nodule in PC69 (Fig. 6AC). A similar transition was observed for *Desulfobacterota* lineages, where the dominance of ANME partners Seep-SRB1 and Seep-SRB2 was succeeded by the persistent, though minor (<10%) presence of *Desulfatiglans* (Fig. 7B). Additional transitions in the archaeal and bacterial communities occurred in the uppermost 3 cm and deepest horizons of the core below 51 cmbsf, with the near seafloor sediments largely devoid of ANME lineages, but supporting a high diversity of *Desulfobacterota* not recovered in deeper horizons, including Seep-SRB4 and Desulfobulbus lineages, while the deepest horizons (51-54 cmbsf) showed an increase in *Halobacteriota* diversity, with the reappearance of members of the ANME-2c and detection of ANME-3, alongside the reoccurrence of Seep-SRB1. Compared to ANME-2c ASVs, the dominant Seep-SRB1 ASVs in the deepest core interval were different from the dominant ASVs in the intermediate depth horizons (3-27 cmbsf), although we observed a few shared Seep-SRB1 ASVs between these regions (Table S11).

Direct comparisons of the carbonate nodules (n=4) with paired sediments across these transitions revealed strong similarities between representative archaeal and bacterial taxa, including major members within *Halobacteriota* and *Desulfobacterota*, down to the ASV level (Fig. 7, Table S11). Beyond these taxa, both Shannon-Wiener and Inverse Simpson indices suggested the sampled nodule bacterial communities were consistently more diverse than their surrounding sediments, though this trend did not extend to the archaeal communities (Table S3).

The long cores recovered from SMM 863 revealed a slightly different community structure from SMM 800-I. Across both SMM 863 cores (LC74, LC66), we observed a clear transition in the dominant ANME and putative sulfate-reducing bacterial groups around 21-24 cmbsf (Fig. 8), aligned with the approximate depth of the SMTZ at this site. Here, members of the *Halobacteriota* above this transition zone were dominated by ANME-2c and known syntrophic partners belonging to the Seep-SRB1 as well as other uncharacterized lineages within the *Desulfosarcinaceae*. We also identified ANME-3 at varying abundances in the upper (<12 cmbsf) sediment in both cores. Below the transition at 21-24 cmbsf, ANME-2c was succeeded by lineages within ANME-1, primarily ANME-1b (up to 75% relative abundance). This transition corresponded with the appearance of the Seep-SRB2 lineage, previously observed to partner with ANME-1 (43, 49, 50). This depth distribution pattern was distinct from the other main ANME syntrophic partner, Seep-SRB1, which was consistently detected above 1% throughout both cores (Fig. 8B). Interestingly, we observed another transition in both cores, beginning at 36-39 cmbsf in LC74 and at 42-45 cmbsf in LC66, where ANME-2c reappeared alongside ANME-3. Here, the lower abundance of ANME-2c at depth was represented by the same dominant ANME-2c ASVs recovered in the upper sediment, while the ANME-3 ASVs at depth largely differed from those recovered near the top of the core (Table S11). Nodules sampled from this transition (n=3) revealed similar ANME-1b and ANME-1a lineages as in the surrounding sediments, down to the ASV level (Fig. 8A, Table S11), but lacked consistent representation from the other minor ANME lineages, including the uncharacterized ANME-1 group (in LC66) and ANME-2c (in both LC66 and LC74). Similarities were notably less apparent among the *Desulfobacterota*, where lineages present in the host sediment were often absent from the nodules (i.e., Seep-SRB1 and Seep-SRB2) and lineages not observed in the immediate sediment horizon were observed in the hosted nodule (e.g., *Desulfatiglans* in LC66).

**Fig. 8.**
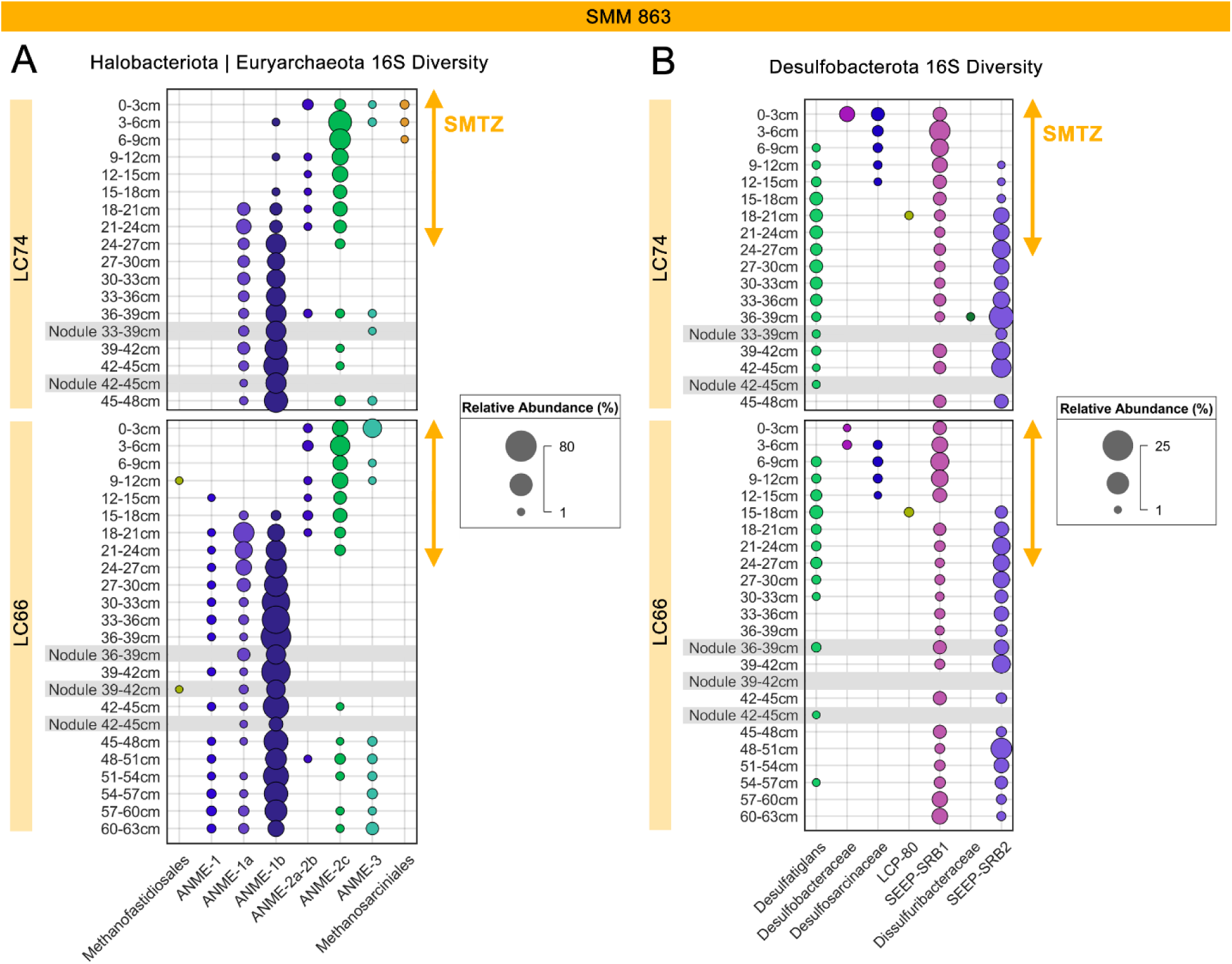
Bubble plots summarizing the depth-resolved, 16S relative abundances for groups within *Halobacteriota*, *Euryarchaeota*, and *Desulfobacterota* in LC74 and LC66 at SMM 863. Where possible, ASVs within these phyla were grouped at the genus-level, or at the next highest taxonomic level possible, and then cut off below a minimum 1% relative abundance. Communities extracted from nodules are highlighted in grey and often represent larger depth horizons. The location of the sulfate-methane transition zone (SMTZ) according to porewater sulfate concentrations from the same or co-located sediment core is indicated with the colored arrows. A) Relative abundances within prominent phyla *Halobacteriota* (featuring ANME) and *Euryarchaeota* (featuring methanogens such as *Methanofastidiosales*). B) Relative abundances within *Desulfobacterota* – the phylum with sulfate-reducing partners to ANME.

### Cell abundance and fluorescence *in situ* hybridization (FISH) in nodules and sediments

Quantification of SYBR Gold®-stained cells and cell aggregates (cell associations > 5 μm) extracts were consistently higher (often 1-2 orders of magnitude) in sediment compared with nodules across the sites, with typical single cell abundances ranging between 10^6^-10^7^ cells mg^-1^ in sediments compared with 10^5^-10^6^ cells mg^-1^ in nodules (Table 3). While differences in cell abundance between sediments and nodules were observed, a substantial decrease in cell abundance between the deeper sediment horizons (e.g., 2.7 x10^7^ at 54-57 cmbsf) compared with shallower depths (e.g., 3.59 x10^7^ at 9-12 cmbsf) was not detected.

**Table 3.**
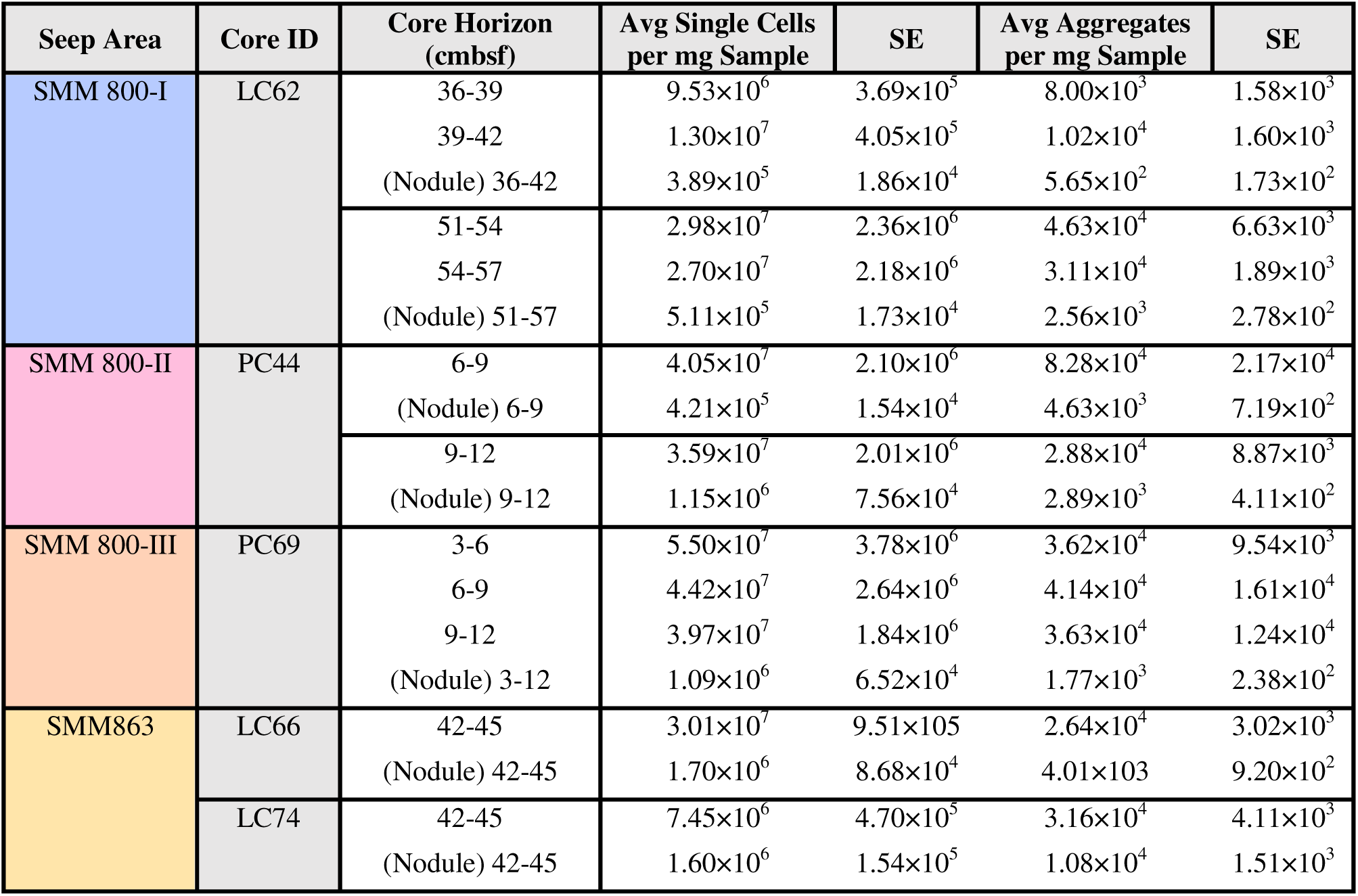
Aggregate/single-cell/cell clump SYBR Gold counts in nodules and paired sediment horizons.

The spatial associations in aggregates and morphology of Bacteria, Archaea, and ANME-1 cells were also examined within the recovered nodules by fluorescence *in situ* hybridization (FISH); (Fig. 9). The persistence of highly auto-fluorescent material within the long cores from seep areas SMM 800-I and SMM 863 made it difficult to clearly discern a positive fluorescence signal from FISH, however we were able to observe positive hybridization with the ANME-1 probe, supporting 16S rRNA results. Notably, we observed large aggregates with morphology consistent with ANME-SRB consortia reported from seep sediments at a higher frequency within nodules recovered from the shallower sediment horizons (< 9-12 cmbsf) at seep areas SMM 800-II and 800-III, compared to the nodules recovered from deeper sediment intervals at SMM 800-I and SMM 863, where the FISH signal from cells was dimmer and cell aggregates were more loosely bound in an exopolymeric matrix (Fig. 9FG, Fig. S14).

**Fig. 9.**
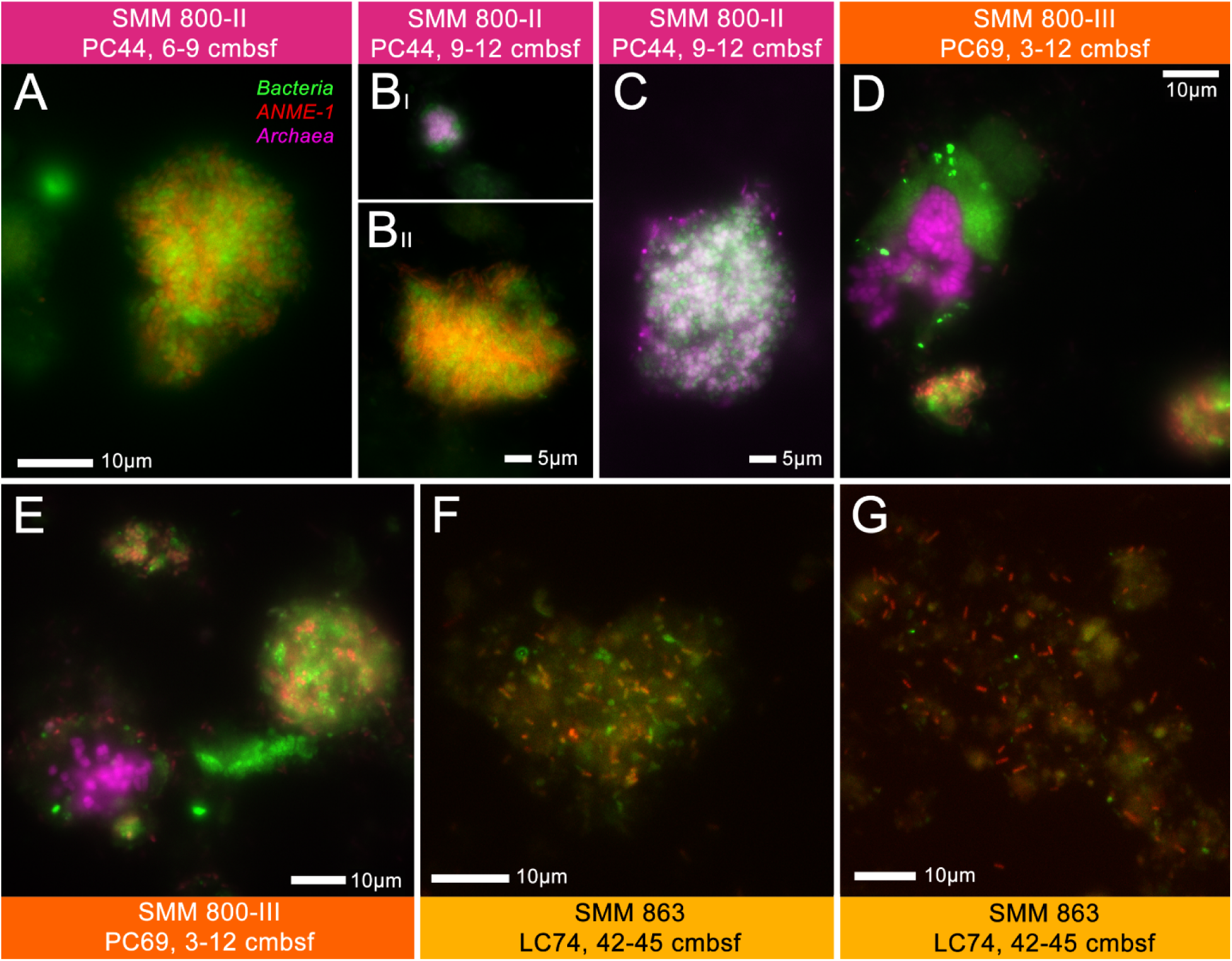
Representative examples of carbonate nodule associated cells and aggregates recovered from SMM 800-II, 800-III, and 863, visualized by fluorescence in situ hybridization (FISH) using probes targeting *Bacteria* (EUB33 I-III, in green), ANME-1 (ANME1-350, ANME1-728; in red), and *Archaea* (ARCH915, in pink). Cell extracts for nodules from SMM 800-I were not shown due to strong interference from auto-fluorescent exopolymeric material.

### Single-cell translational activity measurements (BONCAT) show large shift in nodule endolithic activity between shallow sediments and deeper horizons below the SMTZ

A subset of nodules (n=7) from 5 sediment cores were separated from their host sediment and incubated with the methionine analog HPG under sulfate- and methane-rich conditions to examine translational activity by endolithic microorganisms using biorthogonal noncanonical amino acid tagging (BONCAT), e.g., (49, 51, 52). Nodules were recovered after a 14-week incubation at 4°C, and BONCAT analysis of carbonate-associated cells showed high heterogeneity in activity for both single cells and cell aggregates between nodules, ranging from less than 1% up to 39% of the recovered cells. The relative fraction of extracted cells and cell aggregates (>5 µm) with a confirmed BONCAT signal compared to those without is provided in Table 4. Representative examples of BONCAT stained cells and aggregates from each of the seep areas are shown in Fig. 10.

**Fig. 10.**
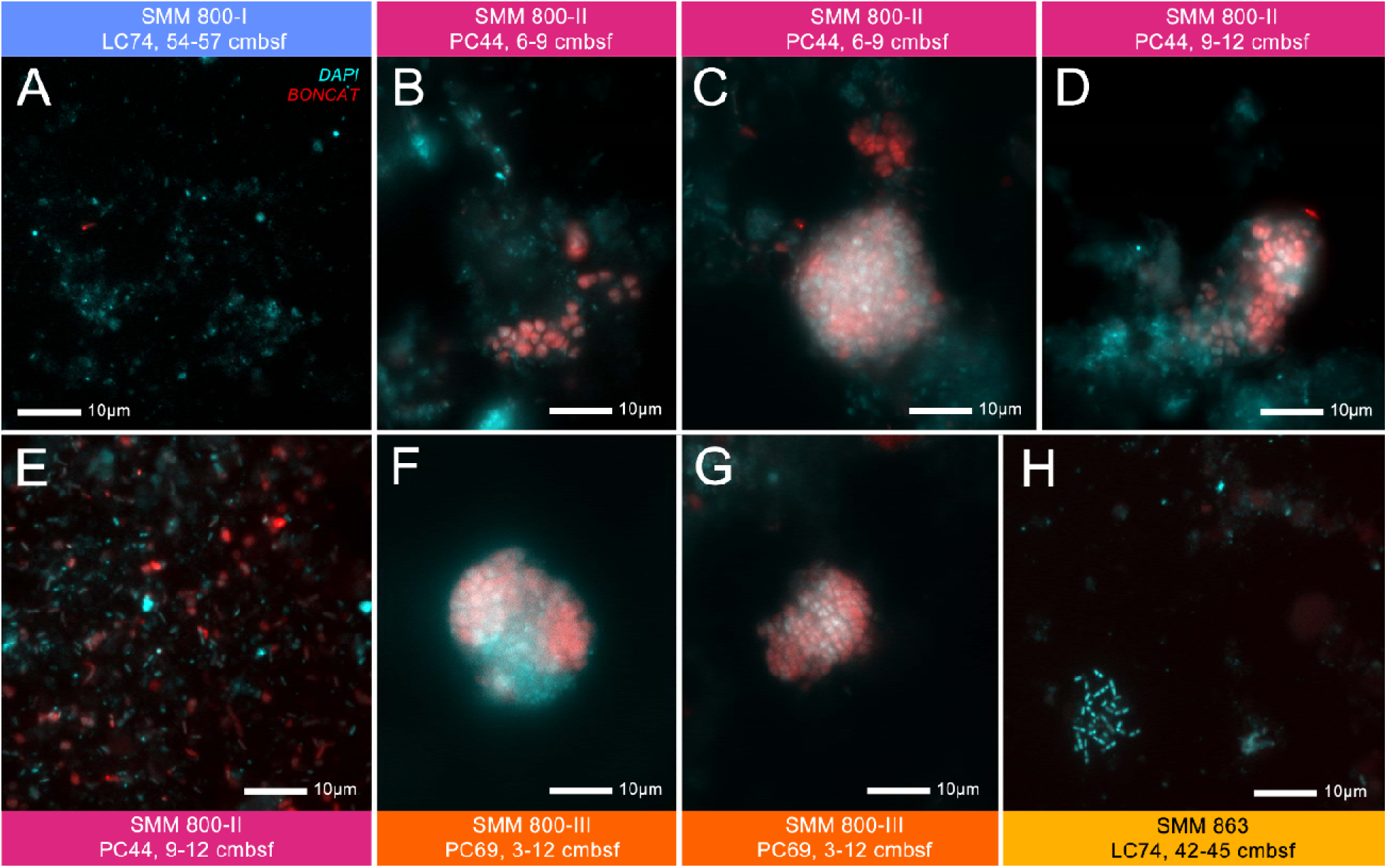
Cell extracts from recovered carbonate nodules at all four seep areas, displaying the presence/absence of newly synthesized proteins with BONCAT (in red) after a 14-week incubation with methionine analog, L-Homopropargylglycine (HPG). DAPI staining of nucleic acids indicating cells is in cyan. Positive BONCAT signal was consistently observed in all extracts from the shallower nodules at SMM 800-II and 800-III, compared to the consistent lack of signal at SMM 800-I and 863.

**Table 4.**
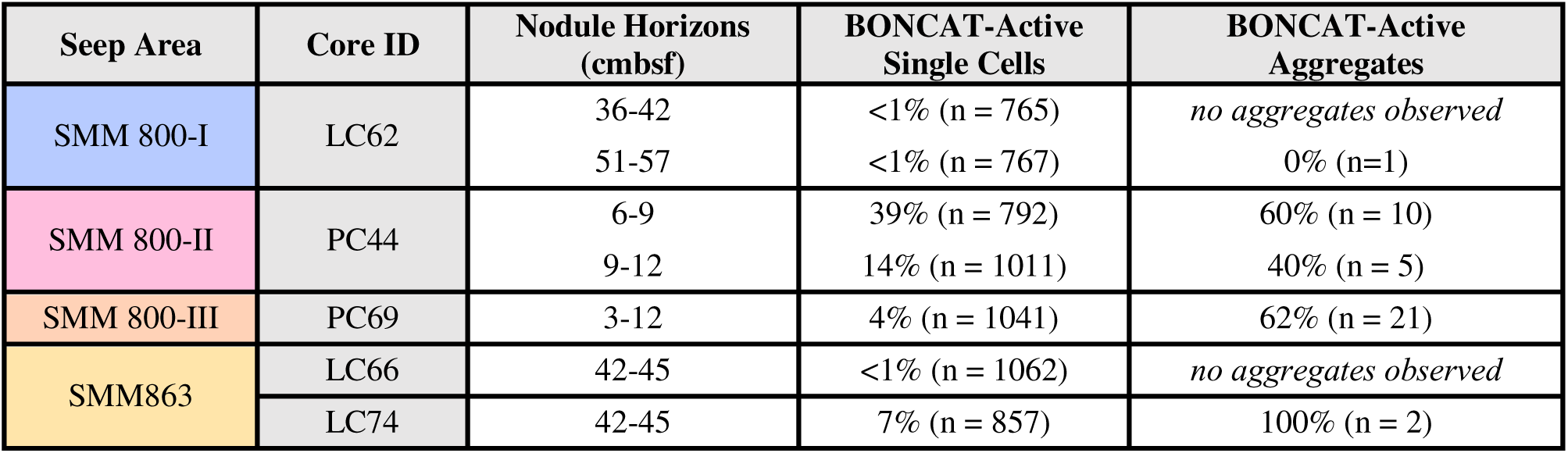
Proportion of BONCAT-active cells to non-BONCAT-active cells from incubated nodules.

| Seep Area | Core ID | Nodule Horizons (cmbfsf) | BONCAT-Active Single Cells | BONCAT-Active Aggregates |
| --- | --- | --- | --- | --- |
| SMM 800-I | LC62 | 36-42 | <1% (n = 765) | *no aggregates observed* |
|  |  | 51-57 | <1% (n = 767) | 0% (n=1) |
| SMM 800-II | PC44 | 6-9 | 39% (n = 792) | 60% (n = 10) |
|  |  | 9-12 | 14% (n = 1011) | 40% (n = 5) |
| SMM 800-III | PC69 | 3-12 | 4% (n = 1041) | 62% (n = 21) |
| SMM863 | LC66 | 42-45 | <1% (n = 1062) | *no aggregates observed* |
|  | LC74 | 42-45 | 7% (n = 857) | 100% (n = 2) |

Nodule-hosted cells recovered from SMM 800-I (LC62, 36-42 and 51-57 cmbsf, both well below the SMTZ), did not exhibit evidence of HPG assimilation among the >700 cells examined per sample. Additionally, these nodules lacked cell aggregates commonly found in shallower sediment horizons. The absence of a BONCAT signal aligns with the minimal sulfide production observed prior to the HPG incubation period (Fig. S15). We observed similar results with a deep nodule recovered from SMM863 below the SMTZ (LC66, 42-45 cmbsf). However, the nodule recovered from the parallel core (LC74) from the same depth horizon did show a weak BONCAT signal (7% of total cells screened) in single cells and aggregates, despite lack of observed sulfide production prior to HPG incubation (Fig. S15). This signal indicated that these endolithic habitats below the SMTZ continued to support viable microorganisms (Table 4).

The low microbial activity and lack of cell aggregates observed in the majority of nodule incubations from the deeper sediment intervals contrasted with the nodules recovered from the shallow horizons within the SMTZ (between 3-12 cm) at SMM 800-II and 800-III. Here, we observed active HPG assimilation for both single cells and cell aggregates, with up to 39% of the single cells (n=792) and 62% of aggregates (n=21) showing a positive BONCAT signal (Table 4). FISH-based identification of BONCAT positive cells indicated a large fraction of these active cells were Archaea (ARCH915 probe, Fig. S16). These samples actively produced sulfide prior to the incubation, reaching values as high as 8 mM after 30 days (Fig. S15), supporting their likely involvement in sulfate-coupled methane oxidation.

## Discussion

In this study, increased depth sampling directly within active seeps using ROV-deployed long cores provided a window into microbial community transitions within and below the SMTZ. We sampled seven sediment cores containing carbonate nodules directly within microbial mat-associated, active seep habitats from four seep areas associated with two carbonate mounds in the Santa Monica Basin. The three seep areas at SMM 800 showed consistent geochemical profiles with high, sulfate-coupled AOM activity characterized by a shallow (<9 cmbsf), sharply defined SMTZ, whereas the two long cores from SMM 863 instead revealed a much deeper (∼21 cmbsf) sulfate depletion zone suggestive of lower methane flux and AOM activity (21). Carbonate nodules recovered from all four seep areas across these distinct geochemical zones enabled us to investigate the physical properties, along with the diversity and metabolic activity of nodule-hosted microbial communities relative to their host sediments, with implications for the degrees to which these incipient carbonate nodules preserve records of past AOM and continue to serve as habitats for *in situ* endolithic communities.

### Closely-linked sediment and nodule-hosted microbial communities indicate ongoing exchange throughout and below the SMTZ

16S rRNA-based community profiles from recovered nodules at all the characterized seep areas reinforced a close similarity to those of their host sediments, regardless of depth and geochemical regime. Non-metric multidimensional scaling (NMDS) ordination of all sediment and nodule-derived ASVs (Fig. S13) also showed a consistent, area-specific clustering between nodules and sediments. This coupling generally agrees with previous studies comparing various carbonate substrates, including nodules, from seep sites at Hydrate Ridge, OR; Eel River Basin, CA; and the Sea of Japan (14–16, 22).

Collectively, these shared microbial community profiles across major geochemical transition zones are strong evidence of continuous colonization and response to *in situ* conditions which collectively influence nodules and sediments, rather than a time-integrated, microbial ‘thumbprint’ of the surrounding sediment.

This is further supported by the high open porosities measured for the nodules in this study, regardless of depth and pore size distributions (< 350 µm), which likely facilitate continuous colonization by surrounding seep sediments. Indeed, 13-month seafloor colonization experiments of sterilized carbonate substrates within and on the periphery of deep-sea methane seeps previously demonstrated active colonization by seep sediment microorganisms, including ANME-1 (16). Although the mechanisms of potential exchange between nodules and sediments remain unknown — especially since motility has not been definitively observed in ANME — previous studies have hypothesized the involvement of fluid advection and/or methane ebullition in facilitating active carbonate colonization (14, 16, 53, 54).

Further, the high nodule open porosities support sufficient connectivity for methane and sulfate perfusion, along with the diffusion of small organic molecules (e.g., HPG, used in BONCAT experiments). As such, these nodules likely experience very similar, current geochemical pressures as the surrounding sediments, which supports the shared microbial community transitions observed in both sediments and nodules from SMM 800 and SMM 863. Indeed, subsurface methane fluxes can vary over periods that can last from days to centuries (55–57), where changes in geochemical stratification exert an active role in structuring the sediment microbial communities performing AOM and facilitating authigenic carbonate precipitation (1, 14, 36, 58–60).

Isotopic and mineral signatures from the nodules recovered below the SMTZ also record carbonate precipitation in different geochemical regimes from the present environments observed. Despite being a sulfate-depleted environment, the consistently depleted δ^13^C values for these nodules — below that of marine organic carbon, -25‰ (61) — support at least a partly AOM-driven origin, and thus preclude exclusive precipitation under the current geochemical conditions. Indeed, all the nodules at SMM 800-I were aragonite-rich (78.2-85.2%), suggestive of a history largely informed by shallower, sulfate-bearing sediments (62, 63). By contrast, the greater Mg-calcite content (45.1-65.1%) in the nodules from SMM 863 support a longer-lived interval in a deeper, more sulfate-limited AOM environment (6, 59, 64, 65). Together, these data suggest that the nodules recovered at depth are likely to have experienced microbial community shifts to reflect communities better adapted to *in situ* conditions, which agrees with a model of continued colonization from the surrounding sediments.

### Persistence of ANME lineages suggest potential adaptation or survival under sulfate-poor conditions below SMTZ

Long cores collected from both SMM 800-I and SMM 863 revealed the consistent dominance of ANME-1 lineages over ANME-2 in the deeper sulfate-depleted sediments and nodules, consistent with several previous studies that have noted this general pattern in seep ecosystems worldwide (24, 26, 28, 30–32, 66, 67). The occurrence of ANME-1 lineages well below the SMTZ, frequently representing the same ASVs as those found within zones of high AOM activity, may reflect thus greater physiological plasticity among this archaeal lineage.

Interestingly, the difference in community profiles below the SMTZ between SMM 800-I (LC62) and SMM 863 (LC74, LC66) attests to additional environmental factors driving niche differentiation in seep sediments and nodules in this geochemical zone. While the dominance of ANME-1b below the SMTZ at SMM 863 may be linked in part to AOM with known ANME partners Seep-SRB1 and Seep-SRB2, the low abundance (<1%) of *Desulfobacterota* sequences below the SMTZ at SMM 800-I suggest that the ANME-1a cells at depth may persist without a sulfate-reducing partner. FISH data from several environments have documented the occurrence of ANME-1 as single cells without an attached bacterial partner (23, 66, 68–71), further supportive of ANME-1’s metabolic flexibility beyond sulfate-coupled AOM.

Notably, the occurrence of putative methylotrophic methanogens within the *Methanofastidiosales* at depth supports the transition to a methanogenic regime below the SMTZ, where ANME-1a and *Caldatribacteriota* (exclusively JS1, Table S11) were found to co-occur. JS1 has been previously identified in anaerobic and/or methanogenic seep sediments worldwide (16, 26, 72–74), and hypothesized to participate in hydrocarbon degradation that may supply key substrates for methanogenesis (44, 73–75). Indeed, it has also been hypothesized that at least some members of the ANME-1 may be capable of methanogenesis (31, 70, 76–80). Although porewater methane concentrations do not conclusively support methane production, previous methane measurements at SMM 800 (33) do support a biological origin for methane, based on the purity and isotopic signature (-70.8‰ δ^13^C). Similarly, the observed rise in porewater DIC and associated δ^13^C in the deeper sediment horizons at SMM 800-I (LC62) may be explained by ^13^C-enriched DIC produced during methanogenesis (81, 82). Enriched ^13^C content associated with an increased dolomite fraction in a deeper nodule from SMM 800-I (LC62, 24-27 cmbsf) also agrees with carbonate formation in historically methanogenic sediment regimes below the SMTZ (6, 64) — a phenomenon additionally observed for a sub-SMTZ nodule recovered from SMM 863 (LC74, 33-39 cmbsf).

Beyond methanogenesis, the observed co-occurrences of ANME-1 lineages, *Methanofastidiosales*, and JS1 may also attest to Fe-coupled AOM below the SMTZ at SMM 800-I. Fe-coupled AOM has been observed in sulfate-depleted marine sediments (29, 83–91), where explicit associations between ANME-1a/1b, JS1, and *Methanofastidiosales* have been statistically supported within seep sediments likely hosting Fe-AOM (29). Further, the detection of ANME-3 in the lower (51-57 cmbsf) horizons of core LC62 at SMM 800-I aligns with observations of ANME-3 in sulfate-depleted marine sediments and microcosms supporting Fe-AOM (83, 85). Ultimately, lack of dissolved iron concentrations for the sediment or discernible iron mineral content in the nodules precludes geochemical support for Fe-AOM at SMM 800-I. Still, past observations of this metabolic regime and its connection to frequently observed microbial taxa in globally-distributed seep sediments below the SMTZ — including sediments and nodules at the seep areas in this study — suggest that the extent of AOM below the SMTZ at active seeps may be under-constrained.

More broadly, the co-occurrence of both ANME/SRB and methanogenic lineages at SMM 800-I may also suggest *in situ* mixing of AOM and methanogenic regimes. These regimes have been hypothesized to co-occur in marine sediments (92) and have been reported for sediments in the Bothnian Sea (87). Significant correlations between ANME clades and methanogen groups from sulfate-depleted sediments at the Okinawa Trough have also suggested a degree of cryptic methane cycling (29). Collectively, these studies underscore the complexity of resolving distinct metabolic regimes in seep sediments.

Interestingly, the weaker-to-negligible BONCAT signal in incubated nodules from below the SMTZ at SMM 800-I and SMM 863 under sulfate and methane-replete conditions indicates that substrate concentrations alone may be insufficient to stimulate measurable AOM activity in nodules at depth over the time period of the incubation (3.5 months). This was despite recovering high cell abundances that included both single cells and aggregate morphologies consistent with ANME (including FISH-detectable ANME-1) in the deeper nodules. Together, these data suggest that measuring metabolic activity for incubated nodules recovered at depth may require longer observation periods, or more sensitive measurements.

In this regard, the minimal activity observed for carbonate nodules below the SMTZ may be compared to that of seafloor carbonates in low-activity seep areas. Indeed, *in situ* experiments transplanting seafloor-exposed carbonates from low to high activity seepage areas by (16) reported shifts towards microbial community structures enriched in lineages involved in AOM and sulfur-based metabolisms after 13 months. Further, sulfide measurements from incubated low activity seafloor carbonates in (11) showed that reactivation of AOM in hosted endolithic communities could take months to years. By contrast, previous ^14^CH_4_ radiotracer measurements of methane oxidation rates from low-activity seep carbonates, incubated under similar conditions, showed detectable, if low methane oxidation (<200 nmol after 7 days) after the introduction of AOM-favorable conditions (8).

Collectively, these data may also point to the understudied nature of metabolic dormancy in seep microbial communities. Previous studies have diverged on interpretation of gene-based profiles of seep sediments and buried seep carbonates, where relict, or ‘fossil’ DNA has been proposed to persist in marine sediments (93–95) and cited in buried seep carbonate crusts (58, 59, 90), such that community profiles are not representative of metabolically viable cells. However, recent studies have suggested that ‘fossil’ DNA negligibly affects taxonomic surveys of environmental samples, especially in marine sediments below the bioturbation zone and even across systems where physical protection of DNA (e.g., seep carbonates) may generate bias (96, 97). As such, we consider 16S rRNA-based nodule community profiles to largely reflect the intact cell assemblage observed in nodules from all depths, with clear variation in metabolic activity with increasing depth below the SMTZ, representing a mixture of dormant and active microorganisms within this endolithic habitat.

Future investigations at methane seeps should therefore seek to better characterize the role of AOM in both the sediment and nodules below the SMTZ, which remains poorly understood. Longer (>1 year), lab-based studies examining the dynamics and range of metabolisms leveraged in these methane-rich, sulfate-poor conditions may shed additional information on factors driving metabolic activity in low-energy seep environments and how sensitive these communities are to the types of geochemical changes characteristic of transient methane seep activity.

### Mechanisms, timescales for sediment community entrapment or exchange in nodules remain underexplored

Although continued exposure and exchange between seep nodules and sediments is strongly supported by the observations presented in this study, the processes or conditions that may act to decouple the nodule environment from its surroundings remain to be more clearly defined. Minor offsets observed between sediment and nodule communities in NMDS ordination of all sediment and nodule-derived ASVs (Fig. S13) may reflect subtle substrate-level differences in community structure. Further, we observed higher community richness in shallow sediments compared to nodules at SMM 800-II and 800-III (Table S3). This trend has also been observed previously in methane seeps at Hydrate Ride, OR (14, 22), and may suggest a local environment more favorable to certain microbial taxa. Indeed, lower cell and cell aggregate abundances in the nodules from SMM 800 and SMM 863 relative to the surrounding sediments may result from niche-specific differences and/or likelihood of some barriers to free exchange between sediments and porous carbonate nodules.

Minor discrepancies between sediment-nodule pairs have also been hypothesized to originate from translocation events that disrupt the original emplacement of incipient seep carbonates. These translocation events act on a local basis and may be driven by sudden gas hydrate decomposition or ebullition events (14). Still, the prevalence of this phenomenon is not known, as the formation and depositional history of seep nodules remains poorly constrained. Sediment-hosted seep carbonates are thought to be continuously subject to burial by continuous sediment transport from nearby coasts, uplift from tectonic movements or gas hydrate accumulation (6, 33, 64), and dissolution from nearby sulfide oxidation and/or aerobic methane oxidation (98–103). In addition to precipitation dynamics, these forces are all expected to shape the duration and nature of exchange between sediments and hosted carbonate nodules in ways that outline future paths of investigation. Although the ages of the nodules in this study remain unconstrained (though loosely estimated at 10^2^-10^3^ years, (33)), the data presented in this study suggest that over the interval since their formation, diagenetic forces have not acted to significantly decouple sediment and nodule microbial communities.

Additional sampling of seep sediment-hosted nodules should seek to more clearly define the time-constrained formation and diagenetic history captured by these understudied carbonates relative to their surrounding sediments, especially in instances where nodule-hosted microbial communities and surrounding sediment communities differ. Uranium-thorium (U-Th) dating, combined with δ^13^C and X-ray diffraction (XRD) characterizations of these sediment-hosted carbonates can better address their age, provenance, and subsequent diagenetic history. In turn, these studies can advance our understanding of controls informing continued microbial habitation in seep sediment-hosted nodules.

## Conclusions

In this study, we integrated microbiological and geochemical analyses of paired seep carbonate nodules and host sediments spanning the SMTZ and below at two previously described seep sites in the Santa Monica Basin, CA. We extend prior characterizations of seep sediment-hosted nodules with this integrated analysis, mapping changes in nodule-hosted microbial community structure and activity of these endolithic microorganisms across steep geochemical and microbiological transitions. By sampling within and below the SMTZ, we captured similar community transitions in both nodules and sediments, suggestive of continued perfusion and exchange between nodules and surrounding sediments, regardless of geochemical conditions. Porosity and mineralogy measurements also indicated sediment-nodule connectivity was feasible and had likely persisted throughout the nodule’s formation, regardless of formation environment.

Unique community structures beneath the SMTZ also attested to different metabolic regimes captured in both sediments and nodules, with the potential for co-occurrence of methanogenesis and AOM. Still, BONCAT activity measurements under sulfate/methane-rich conditions showed that the deeper nodules collected below the SMTZ were slow or no longer capable of producing measurable evidence of AOM within the 14-week incubation period compared to nodules from near-seabed horizons supporting active, sulfate-coupled AOM. Future investigations should explore the range of AOM and non-AOM metabolisms feasible in seep sediment ecosystems in wider detail and seek to more accurately constrain the range of diagenetic histories captured by seep nodules to better constrain the past and current degree of AOM activity recorded in these endolithic habitats.

## Materials and Methods

### Sample collection

Sediment cores were collected with the manipulator arm using the *ROV Doc Ricketts* during dives DR1329 and DR1333 during the WF05-21 Southern California cruise on the *R/V Western Flyer*. Additionally, we included data from one core (PC64) collected on dive DR1248 by ROV *Doc Ricketts* at SMM 800 in February 2020 during the WF02-20 Southern California cruise on the *R/V Western Flyer*. Additional information on site contexts and additional sample recovery details may be found in the Supplemental Material. A further detailed summary of all the sediment and nodules in the study is also included in Table S10.

Upon recovery, all cores were kept in a 4°C walk-in cooler until processing. Within hours of collection, sediment cores were documented and sectioned shipboard into 1- or 3-cm horizons by upward extrusion. Nodules were manually identified during core sectioning and kept within the associated sediment horizons. These horizons were then bagged in sterile Whirl-Pak bags, sealed in Ar-flushed Mylar bags (IMPAK Corporation), and maintained in cold storage at 4°C for, and during, transport to the laboratory.

In the laboratory (<2 weeks after sample collection), nodules >1cm in diameter were manually extracted from the bagged, anoxic sediment horizons under N_2_ and rinsed three times in 0.22 μm-filtered, 3X phosphate-buffered saline (PBS, as in (104)) to remove any adhered sediment. After rinsing, nodules from each horizon were then sub-sampled for various physical, geochemical, microbiological analyses (further detailed by section titled in the Supplemental Material). Due to limited sample available for multiple analyses, some nodules were combined across adjacent sediment horizons and treated as the representation of a larger segment of the sediment core collected (Table S10). We conducted a second cleaning step for the rinsed nodules used for DNA-based community analysis and visualization by fluorescence microscopy to minimize sediment contamination following the optimized protocol for DNA extraction from carbonate nodules described in (14). In (14), rigorous testing for sufficient removal of surface contamination from the sediment hosted nodules was tested using *E. coli* as a tracer, with the following protocol demonstrating complete removal of DNA from the exogenously introduced microbes. Briefly, this surface treatment included additional rinses with cold, 0.22 μm-filtered 3X PBS, sonication, and centrifugation (additional details are provided in the Supplemental Material). Rinsed nodules designated for activity measurements with bioorthogonal non-canonical amino acid tagging (BONCAT) (51) were immediately transferred to separate, sterile, acid-washed media bottles, submerged in cold, 0.22 μm-filtered, N_2_-sparged artificial seawater (media composition available in Supplemental Material Table 1.10), and stoppered with an acid-treated butyl rubber stopper (Th. Geyer GmbH & Co. KG). After immersion, bottle headspaces were flushed with pure CH_4_ for 1 minute with a needle and then pressurized to 2 bar. Samples were incubated in the dark at 4°C.

### Methane measurements

On both the WF05-21 and WF0-20 cruises, sediment methane samples were collected by 1 mL cut off syringe from each sediment core horizon and immediately transferred into 3 mL of 5M NaOH in a pre-weighed gas-tight 10 mL Restek vial, shaken to mix, and then stored upside down at room temperature until analysis. In the laboratory, methane concentrations in the headspace of the vials were measured with a Hewlett Packard (HP) 5890 Series II Plus Gas Chromatography system with a flame ionization detector, connected to a HP 7694 Headspace Sampler with a 1 mL sample loop. The GC column was a 30 m HP-Plot/Q column with inner diameter (I.D.) of 0.32 mm. The software used for analysis was Agilent GC ChemStation (version A.09.03 [1417]). A calibration series ranging from 250 ppm (v/v) to 5000 ppm (v/v) was run immediately prior to samples using lab grade CH_4_ gas (100%) injected into sealed 10 mL Restek vials using a gas-tight syringe. Methane volumetric ppm values were converted to μmol assuming ideal gas behavior at standard temperature and pressure. These values were normalized relative to the mass of sediment added to the Restek vials during the cruise, where masses were corrected for porosity by weighing sediment from each horizon before and after drying in an oven.

### Sulfate measurements

On both cruises, sediment porewater was collected in pressure-driven ‘squeezers’ (105), using argon to compress the sediment section against a 47 mm, 0.22 μm polycarbonate filter (Thermo Fisher) to recover sediment-free porewater in gas-tight, 60 mL syringes while minimizing exposure to air. Porewater from each sediment horizon was then filtered through a 0.22 μm polyethersulfone (PES) syringe filter (Tisch) into 2 mL Eppendorf tubes and stored at -20°C for later analysis. In the laboratory, the concentration of major ion species (e.g. sulfate) was determined by ion chromatography at the Resnick Sustainability Institute’s Water and Environment Lab at the California Institute of Technology. Specifically, samples were run on a 250 mm Dionex IonPac instrument (Thermo Fisher) with AS19-4μm and CS16-4μm columns for anions and cations, respectively — each with 50 mm versions as guard columns. 100 μL of each porewater sample were diluted 50x in milliQ ultrapure DI water (final volume 5 mL) before running. Standard curves were generated using calibration standards diluted similarly to samples, with 100 μL of 500 mM NaCl added to mimic typical seawater sample peak behavior. Analyte peaks were automatically integrated and major ion species, including sulfate, were automatically calculated using the prepared standards with the Chromeleon Dionex IC (Thermo Fisher, v7.2.9) software, with manual inspection to ensure regular peak shapes.

### Sulfide measurements

Samples for porewater sulfide concentrations were prepared by the addition of 0.5mL of 0.22 μm filtered porewater from each sediment horizon to 0.5 mL of 0.5M zinc acetate in a 1.5 mL Eppendorf tube to precipitate zinc sulfide. Due to its lability, sulfide samples were processed first after porewater collection from the squeezer apparatus. In the laboratory, the precipitated sulfide was quantified spectrophotometrically according to the Cline sulfide assay protocol (106). Briefly, this involved the acidification of the sample and reaction with N, N-dimethyl-p-phenylenediamine dihydrochloride in presence of ferric chloride. The resulting methylene blue solution’s absorbance was measured in duplicate at 670-nm wavelength with a microplate reader (Sunrise, Tecan). Concentrations were calculated after subtracting absorbance values from DI H_2_O blanks from concurrently measured zinc sulfide standards prepared from sodium sulfide solutions (0.12 mM to 25 mM). We additionally measured 10x and 50x dilutions to allow for quantification of highly sulfidic samples within the linear color range.

### DIC and **δ**^13^C analyses

We measured dissolved inorganic carbon (DIC) concentrations and its δ^13^C in porewater (C_pw_), as well as the δ^13^C of nodule (C_nod_) where sufficient porewater or nodule material allowed. After extraction and rinsing, nodule samples were air dried and ground to a fine powder with an agate mortar and pestle, which was cleaned with DI water and ethanol between samples. Replicate subsamples of powdered nodules (0.45-0.65-mg) were added to 12 mL, acid-washed and combusted Exetainer vials (Labco). The vials were then flushed with helium for 2 minutes and acidified with 200 μL 43.5% phosphoric acid. To measure porewater DIC and δ^13^C_pw_, 1 mL of extracted porewater was immediately added via syringe to 12 mL, acid-washed and combusted Exetainer vials (Labco) pre-flushed with helium and pre-loaded with 100 μL 43.5% phosphoric acid during the WF05-21 and WF02-20 cruises. In the laboratory, the released CO_2_ gas was measured on a GasBench II coupled to a Delta V Plus isotope ratio mass spectrometer (IRMS) at the Stable Isotope Facility at the California Institute of Technology.

Concentrations of DIC were determined based on comparison of the average total peak area of sample injections to replicate sodium bicarbonate standard curves. The 6-point dilution standard curves for the porewater DIC used ultrapure DI H_2_O as a background standard and a solution series (1, 2.5, 5, 10, 20, and 30mM) of sodium bicarbonate (-2.9‰).

δ^13^C_nod_ and δ^13^C_pw_ values were corrected for instrument dependency on CO_2_ concentration and then normalized to the VPDB scale with a two-point calibration (107) using the Merck (δ^13^C = - 49.2‰) and Carrara Marble (CM 2013, δ^13^C = +2‰) carbonate standards. Here, we use solid carbonate powder isotope standards for both δ^13^C_nod_ and δ^13^C_pw_ on the basis of a quantitative conversion (i.e., total carbonate consumption) and since O isotopes are not considered for this study.

### Nodule porosity & porosimetry

During nodule sub-sampling, select nodules (where sufficient material allowed) were additionally rinsed in deionized water to remove any salts and dried at room temperature for structural and mineralogical analysis. To characterize the internal structure of the recovered carbonate nodules, we determined their open porosity using a water intrusion (or Archimedes) method. We followed Method A of ASTM D792 (108), which collects the dry mass, the immersed mass (i.e., mass of fluid displaced), and the saturated mass (i.e., mass of fluid-saturated object) of a nodule sample with DI water, according to Eq. 1:

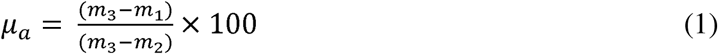

where *μ_a_* is the apparent, or open, porosity (expressed as a percentage of the sample bulk volume), *m_1_* is the dry mass, *m_2_* is the immersed mass, and *m_3_* is the saturated mass. This was achieved with a density apparatus attached to a fine scale (Mettler Toledo). Additionally, we measured the pore size distribution of paired samples with Hg-porosimetry. This method uses a pressurized, non-wetting liquid (mercury) to intrude the nodule pore spaces, where the cumulative volume of intruded mercury is recorded at each pressure interval, corresponding to a certain pore size (109). We used a Micromeritics AutoPore IV Mercury Intrusion Porosimeter to measure open pore size distribution between 6 and 350 µm by recording cumulative mercury intrusion volume at 90 intervals within a pressure range of 3.51 and 206 kPa.

### XRD

Nodule subsamples for mineralogical characterization with X-ray diffraction (XRD) were air-dried and ground to fine powder with an agate mortar and pestle, cleaned with ethanol between each use. Resulting powders were analyzed with a Rigaku SmartLab powder diffractometer at the X-Ray Crystallography Facility in the Beckman Institute at the California Institute of Technology. We used a Cu Kα source at 50 kV and 40 mA. Scans were run from 5° to 70° 2θ at a scanning speed of 0.02° 2θ/s. Rigaku SmartLab Studio II was used to fit and identify peaks and the 2020 Crystallography Open Database was used for phase identification. The relative semi-quantification of clearly identifiable phases was calculated according to the reference intensity ratio (RIR) based on a corundum standard and documented in weight percent. For the phases, the RIR method used the (104) peak heights of calcite, Mg-calcite and dolomite, the (111) peak height of aragonite, and the (101) peak height for quartz. Here, Mg-calcite corresponds to (Ca_0.9_Mg_0.1_)O_3_, as measured by (35).

### 16S rRNA gene sequencing

To characterize and compare the diversity and composition of the microbial communities across the seep areas represented in the dataset, we sequenced the V4 and V5 regions of the 16S rRNA gene. Here, 2mL of sediment from each core horizon was sampled by sterile 1 mL cutoff disposable syringe into a 2 mL screwcap Eppendorf tube and immediately stored at -80°C for DNA extraction. In the laboratory, doubly rinsed nodules intended for sequencing were also finely ground and frozen at -80°C after extraction from sediment and cleaning as described above. DNA was extracted from 0.15-0.3 g (wet weight) of thawed sediment or finely ground nodule powder using the DNeasy PowerSoil Pro Kit (Qiagen). Target sequences were then amplified with a 2-step Illumina sequencing strategy (110). Briefly, this involved an initial amplification with 515f (5’-TCGTCGGCAGCGTCAGATGTGTATAAGAGACAG-GTGYCAGCMGCCGCGGTAA-3’) and 926r (5’-GTCTCGTGGGCTCGGAGATGTGTATAAGAGACAG-CCGYCAATTYMTTTRAGTTT-3’) primers, followed by a second amplification to include the necessary Illumina barcodes prior to being sequenced.

Adapters from paired-end sequences were removed using ‘Cutadapt’ (v2.9), (111). Trimmed reads were then input into QIIME2 (v2020.11), (112), where we used a DADA2 pipeline (v1.14.1), (113) to filter and denoise the reads as well as remove chimeric sequences. ASVs with assigned taxonomy were generated from the merged, non-chimeric sequences in QIIME2 using the Silva V138.1 SSURef database feature classifier (114). Classified, non-zero ASVs for all the sediment and nodules in the study are included in Table S11. Additional details on modified DNA extraction protocols, PCR amplification, gene sequencing, and read processing are also included in the Supplemental Material.

During subsequent analysis, we calculated Shannon-Wiener and Inverse Simpson diversity indices for archaeal and bacterial ASVs from the carbonate nodules and their host sediments. Further information on how these indices were calculated are detailed in the Supplemental Material. Additionally, we performed a non-metric multidimensional scaling (NMDS) ordination of all carbonate nodule and sediment-derived ASVs. Further details on how this ordination was performed is also detailed in the Supplemental Material.

### Fluorescence in situ hybridization (FISH)

During sediment core sectioning, 0.5-0.7 mL sediment per horizon was immediately added to 0.5 mL of a 0.22 μm filtered 4% paraformaldehyde solution in 3X PBS and fixed overnight at 4°C. Afterwards, the sample was washed twice with 0.22 μm filtered 1X PBS and then re-suspended in a 50% ethanol: 1X PBS medium and stored at -20°C. During nodule extraction in the laboratory, doubly rinsed nodules were finely ground with a sterile agate mortar and pestle.

Immediately after grinding, the ground nodules were also fixed and washed as was done for the sediments, but re-suspended in a 70% ethanol: 1X PBS medium prior to storage at -20°C. To visualize the cells with minimal interference from sediment and mineral particles, we then performed a density separation using percoll, based on a modified protocol outlined in (8, 23, 115). Briefly, 100-200 μL fixed sediment or carbonate was added to 400 μL of a chilled, 0.22 μm filtered ‘combination’ buffer made of 0.5% v/v Tween 20, 3 mM sodium pyrophosphate, and 0.35% wt/v PVP (116). Samples were then sonicated on ice over 3 cycles at amplitude 20 on a Q55 sonicator (Qsonica), with 10 seconds of active sonication and 10 seconds of rest in between to avoid overheating. Next, sample tubes were shaken on a vortex mixer (Vortex-Genie 2, Scientific Industries) for 30 minutes at room temperature. We then added 400 μL of chilled, 0.22 μm filtered 1X PBS before mixing and overlaying the mixed sample on 950 μL of a chilled percoll density gradient. The gradient tubes were then centrifuged at 18,000 x g for 25 minutes at 4°C (Microfuge 18, Beckman Coulter). After centrifugation, the density gradient supernatant was transferred to a new set of sample tubes and centrifuged again at 16,000 x g for 3 minutes at 4°C to form cell pellets. The pellets were washed twice with 0.22 μm filtered 1X PBS.

The washed cells were then stained with a SYBR® Gold (Thermo Fisher) solution at a 25X final concentration in ultrapure DI H_2_O in the dark for 15 minutes at room temperature. After staining, the cells were centrifuged at 16,000 x g for 3 minutes at room temperature, re-suspended in ultrapure DI H_2_O, and stored overnight at 4°C. The next day, the cells were mounted via vacuum filtration onto a 0.2 μm black polycarbonate filter and a 5μm PVDF backing filter (Millipore Sigma). The filters were washed twice in ultrapure DI H_2_O, air dried, and then stored in the dark at 4°C until counting. Prior to coverslip placement, each filter received ∼ 50 μL Vectashield® Antifade Mounting Medium (VectorLabs). For counts, we used an Olympus BX51 fluorescence microscope using a 100x objective lens (Olympus). Thirty fields of view per filtered sample were surveyed to determine the number of cell aggregates and single cells. For these counts, any association of cells larger than 5 μm were considered aggregates. Cell and aggregate counts were normalized per unit dry mass for all sample types.

To observe the presence and morphologies associated with particular taxa, we used fluorescence in situ hybridization (FISH). For FISH, the washed, percoll-extracted cells were re-suspended in ultrapure DI H_2_O and stored overnight at 4°C. The following day, cell suspensions were dried at room temperature onto separate 6 mm wells on PTFE- and L-lysine-coated glass slides (Tekdon). The hybridization probes EUB338 [5’-GCTGCCTCCCGTAGGAGT-3’ (117), 3’ conjugated Alexa Fluor™ 488 dye], EUB338 II [5’-GCAGCCACCCGTAGGTGT-3’ (118), dual-labeled Alexa Fluor™ 488 dye], EUB338 III [5’-GCTGCCACCCGTAGGTGT-3’ (118), dual-labeled Alexa Fluor™ 488 dye], ANME1-350 [5’-AGTTTTCGCGCCTGATGC-3’ (19), dual-labeled Alexa Fluor™ 546 dye], ANME1-728 [5’-GGTCTGGTCAGACGCCTT-3’, designed using ARB (119) to hit the ANME1 Clade of the Silva 138.1 database), 5’ conjugated Alexa Fluor™ 546 dye], and ARCH-915 [5’-GTGCTCCCCCGCCAATTCCT-3’ (120), dual-labeled Alexa Fluor™ 647 dye] were used for the mounted cell extracts. The hybridization probe mix was prepared in sterile, 40% formamide concentration, with final concentrations of 5-ng/μL for each probe (Integrated DNA Technologies). Each well on the slide then received 10 μL of probe mix and was hybridized in the dark, overnight at 46°C. The next day, the glass slides with hybridized cells were immersed and incubated in a pre-warmed hybridization wash in the dark for 10 minutes at 48°C. The slides were then rinsed in ultrapure DI H_2_O and allowed to air dry (in the dark) completely. FISH hybridization probe and wash recipes are provided in Tables S4-S5.

Prior to coverslip placement, each well also received 10 μL of a 90% Citifluor^TM^ mounting medium (Electron Microscopy Sciences) with 4.5 ng/μL 4’,6-diamidino-2-phenylindole (DAPI) for nucleic acid visualization. Hybridized cells were then visualized with an ELYRA S.1 / Axio Observer Z1 super resolution microscope using a 100x objective lens (Zeiss). For control experiments, each glass slide had a well with mounted cells subjected to the entire FISH protocol but used ultrapure DI H_2_O instead of probes. For these wells, negligible signal was observed.

### Single cell translational activity measurements with BONCAT

Translational activity was observed by amending a subset of the bottled nodules with an anoxic L-Homopropargylglycine (HPG) solution. HPG works as a methionine analog and is incorporated during protein synthesis without significantly altering community composition or metabolic activity (49, 51, 52). After incorporation, proteins containing HPG can be fluorescently labeled through copper-catalyzed azide-alkyne ‘click’ chemistry, which enables detection via fluorescence microscopy (49, 52, 121, 122).

In this study, the nodules previously maintained in N_2_-sparged artificial seawater with a 2 bar CH_4_ headspace were separated into replicate experimental and control bottles. Where possible, the nodules were selected to minimize mass differences between experimental and control groups. Each new sterile, acid-washed bottle received 50 mL of 0.22 μm-filtered, N_2_-sparged artificial seawater of the same composition (Table S6, sufficient to immerse the nodules).

Throughout transfer and immersion, all bottles were constantly maintained under active N_2_ flow to minimize oxygen exposure. After immersion, all bottle headspaces were flushed with pure CH_4_ for 1 minute and pressurized to 2 bar. Samples were then incubated in the dark at 4°C for a period of 5 weeks, during which the media was periodically sampled to monitor sulfide concentrations in control and experimental bottles. After determining comparable sulfide production between each control and experimental group (Fig. S15), the artificial seawater media was replaced with freshly prepared 0.22 μm-filtered, N_2_-sparged artificial seawater. Additionally, each experimental bottle received aqueous 0.22 μm-filtered, N_2_-sparged HPG to a final concentration of 200 µM. Throughout media replacement and HPG addition, all bottles were constantly maintained under active N_2_ flow to minimize oxygen exposure. After immersion, all bottle headspaces were again flushed with pure CH_4_ for 1 minute and pressurized to 2 bar. Samples were then incubated in the dark at 4°C for a period of 14 weeks to ensure sufficient HPG assimilation.

At the end of the incubation period, nodules from both control and experimental bottles were ground, fixed, and washed as previously described for FISH hybridization. Cells from each fixed, ground nodule were then extracted and mounted onto separate 6 mm wells on PTFE- and L-lysine-coated glass slides as previously described. For the BONCAT dye click reaction, the glass-mounted cell extracts were then dehydrated by sequential immersion in 50% (1 minute), 80% (3 minutes), and 90% (3 minutes) ethanol: DI H_2_O baths and dried room temperature.

We used BONCAT azide dye Oregon Green 488 (VectorLabs) at a final concentration of 20 µM. 10-µL of the dye solution was then added to each well and incubated in the dark for 30 minutes. For the incubation, we flushed a lidded sample container with Ar, in which we placed the mounted cells to minimize oxygen exposure, which can reduce the fluorescent signal. After incubation, the slides were washed in 0.22 μm-filtered 1X PBS for 10 minutes (in the dark), quickly rinsed in ultrapure DI H_2_O, and allowed to air dry in the dark prior to FISH hybridization. Hybridization probes EUB338 [5’ conjugated cyanine 3 dye], EUB338 II [3’ conjugated cyanine 3 dye], EUB338 III [5’ conjugated cyanine 3 dye], and ARCH-915 [dual-labeled Alexa Fluor™ 647 dye] were used. Hybridization was done at 35% formamide. Washing and addition of DAPI-Citifluor® mounting medium was performed as previously described. BONCAT click reaction and FISH hybridization recipes are listed in Tables S7-S9.

Click-dyed, hybridized cells were then visualized with an ELYRA S.1 / Axio Observer Z1 super resolution microscope using a 100x objective lens (Zeiss). Fraction of cells with HPG signal were evaluated by counting HPG-incubated, extracted cells with observable signal above a mean background threshold calculated from the control cells’ auto-fluorescence in the same channel. For FISH and BONCAT control experiments, each glass slide had a well with mounted, non-HPG labeled cells subjected to the entire FISH protocol but used ultrapure DI H_2_O instead of probes. For these cells in these negative control wells, a negligible signal was observed.

## Data availability

The Santa Monica Mounds Low Altitude Survey System (LASS) survey was done in the Spring of 2018 using *ROV Ventana* on *R/V Rachel Carson*. The Santa Monica Mounds Mapping AUV surveys were done multiple times from *R/V Zephyr* and *R/V Carson*. Featured grids and photomosaics will be available at MGDS https://www.marine-geo.org/.

The raw DNA sequence reads generated from the SMM 800 sediment cores in this study were initially submitted to NCBI under project accession number PRJNA1196099 (sample accession numbers: SAMN45814414-SAMN45814434; SAMN45814441-SAMN45814448) and are accessible at https://www.ncbi.nlm.nih.gov/bioproject/ PRJNA1196099. The rest of the raw DNA sequence reads generated for this study (SMM 800 nodules, SMM 863 samples, experimental controls) were submitted to NCBI under accession numbers PRJNA1207588 and accessible at https://www.ncbi.nlm.nih.gov/bioproject/PRJNA1207588.

## Acknowledgements

We are indebted to the captain, crew, and pilots of the *ROV Doc Ricketts* from the WF05-21 cruise, who made this work possible. We also thank David Caress, Charles Paull, Eric Martin, and the Autonomous Systems Operations group at the Monterey Bay Aquarium Research Institute (MBARI), as well as the crews of *R/V Zephyr* and *R/V Carson* for collection, processing, and analysis of bathymetry maps of the Santa Monica Basin. We thank S. Lim (University of Nevada, Las Vegas), J.S. Magyar (Caltech), and S. Goffredi (Occidental College) for their help in sample collection and processing during the WF05-21 Southern California cruise of the *R/V Western Flyer*, S. Connon (Caltech) for technical support and barcoding of 16S amplicons for sequencing, D. Utter (Caltech) and H. Sapers (Northern Arizona University) for support during data processing.

S.A. Parra and V.J. Orphan conceptualized the study and acquired financial support for the study. J. Mullahoo analyzed dissolved porewater methane. L.K. Quinn analyzed nodule open porosity and Hg-porosimetry. R.L. Wipfler analyzed porewater sulfate concentrations. S.A. Parra processed and analyzed all other data presented in the study, including nodule selection, porewater sulfide measurements, DIC/δ^13^C measurements, x-ray diffraction, DNA extraction, 16S rRNA sequence analyses, fluorescence microscopy, and BONCAT incubations. M.J. Mayr and V.J. Orphan supervised the study. S.A. Parra wrote the original draft and incorporated edits before and during the publication stage. M.J. Mayr, J. Mullahoo, L.K. Quinn, R.L. Wipfler, and V.J. Orphan all participated in critical review and commentary on the paper draft before and during the publication stage.

This work was supported in part by NASA’s Interdisciplinary Consortia for Astrobiology Research (ICAR) program under Award Number AWD-005316-G4 (to V.J. Orphan), the National Science Foundation under Award Number 2048666 (to V.J. Orphan), and NASA’s Future Investigators in NASA Earth and Space Science and Technology (FINESST) program under Award Number 80NSSC22K1336 (to S.A. Parra). This material is based upon work supported by the U.S. Department of Energy, Office of Science Biological and Environmental Research Program under Award Number DE-SC0022991 (to V.J. Orphan).

Disclaimer: This report was prepared as an account of work sponsored by an agency of the United States Government. Neither the United States Government nor any agency thereof, nor any of their employees, makes any warranty, express or implied, or assumes any legal liability or responsibility for the accuracy, completeness, or usefulness of any information, apparatus, product, or process disclosed, or represents that its use would not infringe privately owned rights. Reference herein to any specific commercial product, process, or service by trade name, trademark, manufacturer, or otherwise does not necessarily constitute or imply its endorsement, recommendation, or favoring by the United States Government or any agency thereof. The views and opinions of authors expressed herein do not necessarily state or reflect those of the United States Government or any agency thereof.

## Notes

### Competing Interest Statement

The authors have declared no competing interest.

### Summary of Updates

Supplemental Data has been added to the initial submission.

