## Supplemental Material for "Diversity and single-cell activity of endolithic microbes in sediment-hosted carbonate nodules within and below the sulfate-methane transition zone"

Sergio A. Parra[a*]#, Magdalena J. Mayr[a,b], James Mullahoo[a], Laura K. Quinn[c&], Rebecca L. Wipfler[a], Victoria J. Orphan[a,b]#

[a]Division of Geological and Planetary Sciences, California Institute of Technology, Pasadena, California, USA

[b]Division of Biology and Biological Engineering, California Institute of Technology, Pasadena, California, USA

[c]Division of Chemistry and Chemical Engineering, California Institute of Technology, Pasadena, California, USA

Running Head: **Endolithic microbes in carbonate nodules**

[*]Present Address: Sergio A. Parra, Boston Consulting Group, Los Angeles, CA, USA.

[&]Present Address: Laura K. Quinn, Saint-Gobain Ceramics & Plastics, Inc., Northborough, MA, USA

### **Site Descriptions and Sample Collection**

#### *Site Classification and Sample Collection*

At all sampling locations, we confirmed local methane seep activity with in-situ observation of white or orange microbial mats, clam beds, and/or gas bubbling from the seafloor; this classification scheme is consistent with previous biological and geological surveys of seeps around the world (1–3). Sediment push cores (~7.6 cm outer diameter, ~7 cm inner diameter) were collected as two core types: a shorter push core (PC, ~30 cm long) and long core (LC, ~1.22 m long).

#### *SMM 800-I*

SMM 800-I (33.799447N, 118.647171W) is located at an outcrop of seep carbonate on the western flank of the SMM 800 (NE) mound. This site was characterized by a light peach colored microbial mat adjacent to a white mat, shown in Fig. S1.  At this site, the sediment was sufficiently deep to insert 1.22 m long cores with the ROV manipulator. LC62 was taken directly within the light peach mat, with full insertion into the sediment. Upon recovery, we observed bubble textures throughout the core, with large void spaces at 30-cmbsf.

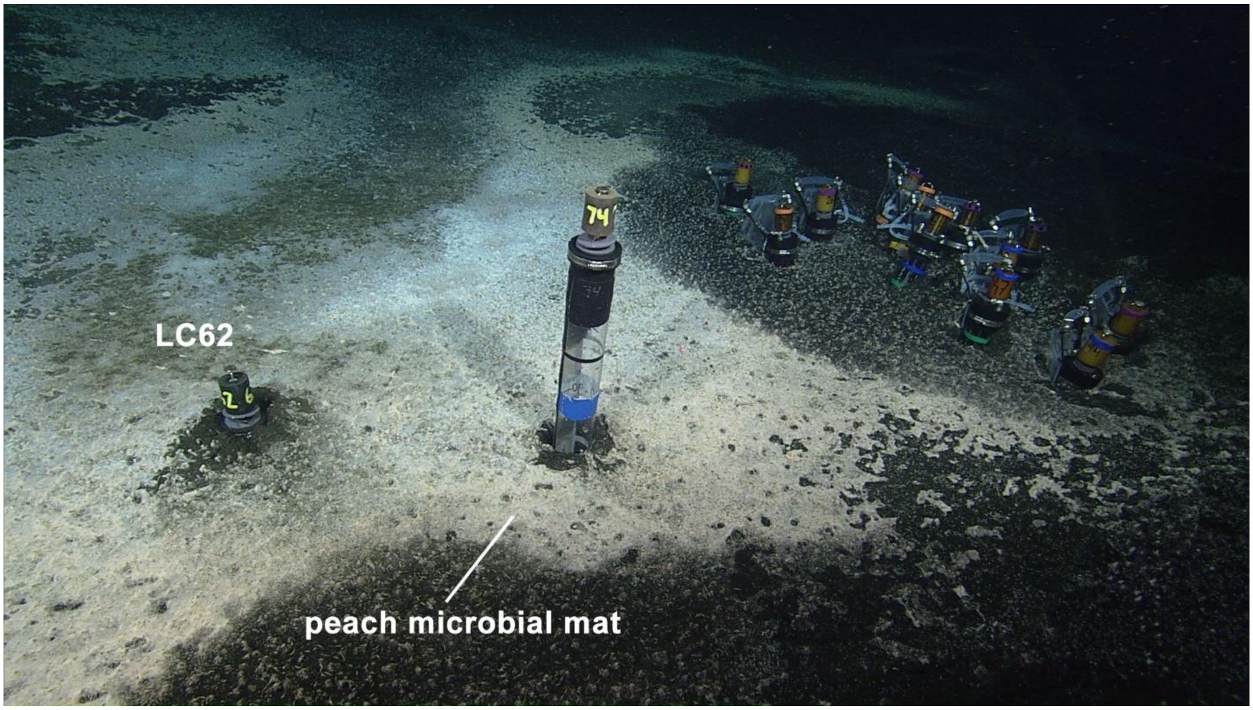


**Fig. S1** – In situ image of the sampling site at SMM 800-I during the WF05-21 cruise. LC62 was sampled over a microbial mat (shown, left). Additional cores were taken (shown, right) but not used for this study. Core diameter is ~7 centimeters.

*SMM 800-II*

SMM 800-II (33.799662N, 118.646947W) is located on the northern flank of the primary SMM 800 mound, about 1 meter down the mound from SMM 800-I. We observed white and thin, grey microbial mats and small clams, shown in Fig. S2A. An initial push coring attempt on the white microbial mat region failed due to a hard layer observed at ~8-10 cmbsf, presumed to be carbonate platform. A second, successful coring attempt was made on the sparser, grey microbial mat. Due to this success, PC44 was taken directly on top of the grey mat, with full insertion into the sediment. The site's appearance in 2021 was notably different than when it was previously sampled in 2020. During the 02-20 Southern California oceanographic expedition of the *R/V Western Flyer*, we observed thick, white and peach-colored microbial mats, which suggested a previously higher degree of methane seepage activity. PC64 was taken on the white microbial mat adjacent to the peach-colored mat, shown in Fig. S2B. The full length (30 cm) was not recovered due to encountering a hard layer at 9 cmbsf (presumed to be carbonate platform), though active bubbling was observed prior to sectioning.

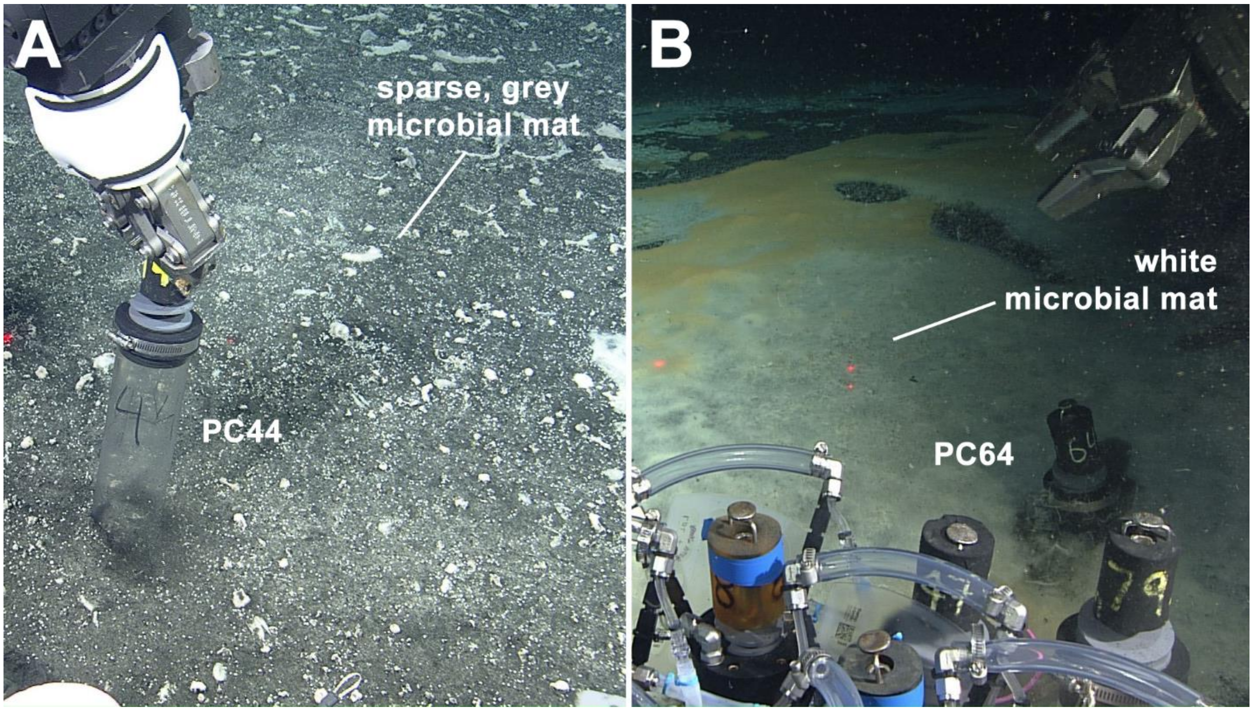


**Fig. S2** – *In situ* images of the sampling site at SMM 800-II. A) PC44 was sampled over a sparse microbial mat during the WF05-21 cruise. B) In contrast, PC64 was sampled over a thicker microbial mat during the WF02-20 cruise at the same site, suggesting a decrease in the degree of methane seepage activity. Core diameters shown are ~7 centimeters. Laser dots in the images are 29 cm apart.

*SMM 800-III*

SMM 800-III (33.798962N, -118.646264W) is located on the southeastern flank of the primary SMM 800 mound, about 5 meters down the mound from SMM 800-II. We observed a large white microbial mat with a patch of dark sediment in the center. Two cores, PC43 and PC69, were taken directly over the white mat on the eastern end of the mat patch, shown in Fig. S3. However, both cores were only half-length cores due to solid carbonate encountered at 9 and 12 cmbsf for PC43 and PC69, respectively. Upon recovery, we observed small white clams at the surface, as well as active bubbling coming from deeper within PC69. For PC43, we observed denser white microbial mat filaments in the top 3-4 cmbsf, and active bubbling as well as trapped gas voids below 8 cmbsf.

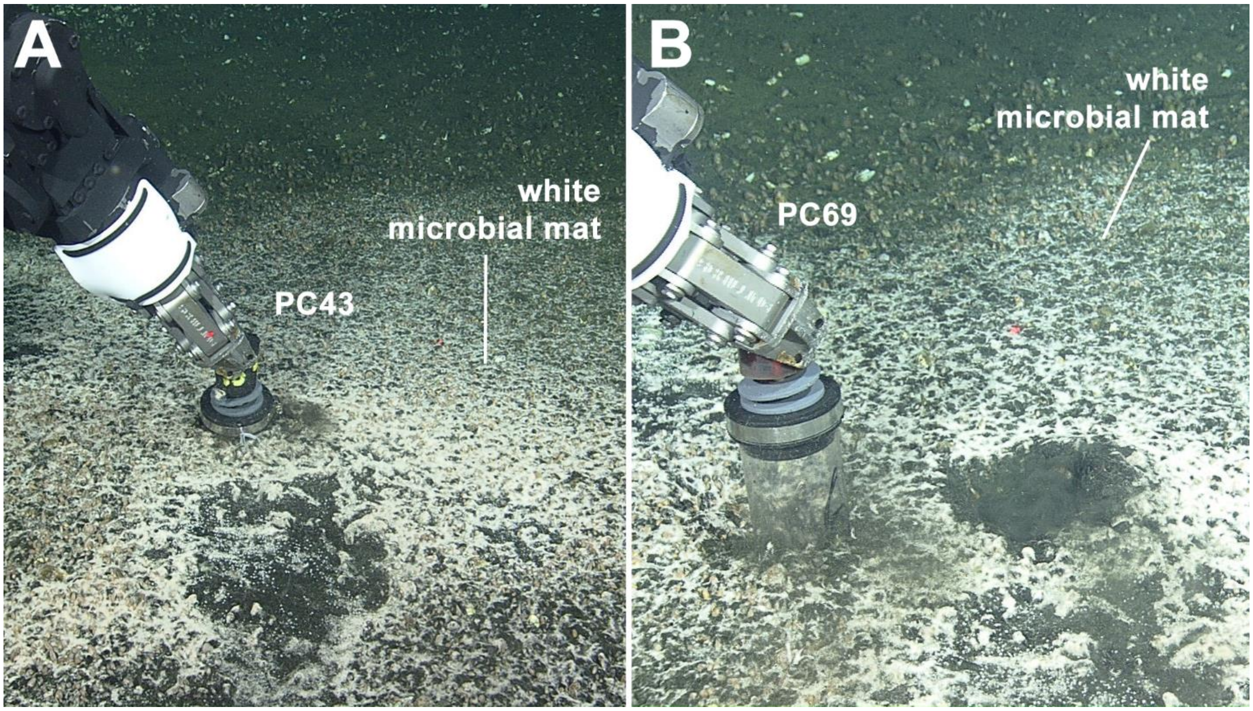


**Fig. S3** – *In situ* images of the sampling site at SMM 800-III during the WF05-21 cruise. A) PC43 was sampled over a white microbial mat. B) PC69 was sampled to the left of PC43 (dark sediment spot to the right) over the same white microbial mat. Core diameters shown are ~7 centimeters. Laser dots in the images are 29 cm apart.

*SMM 863*

SMM 863 (33.788977N, -118.647171W) is located on the southern flank of the primary mound at Site 863 (863m depth), about 2.25km SW of SMM 800. We observed orange and white microbial mats, shown in Fig. S4. LC66 and LC74 were taken directly over the orange mat. Both cores were only able to be partially inserted due to a hard substrate (presumed to be carbonate pavement) at depth, 50-cmbsf for LC74, and 65-cmbsf for LC66.

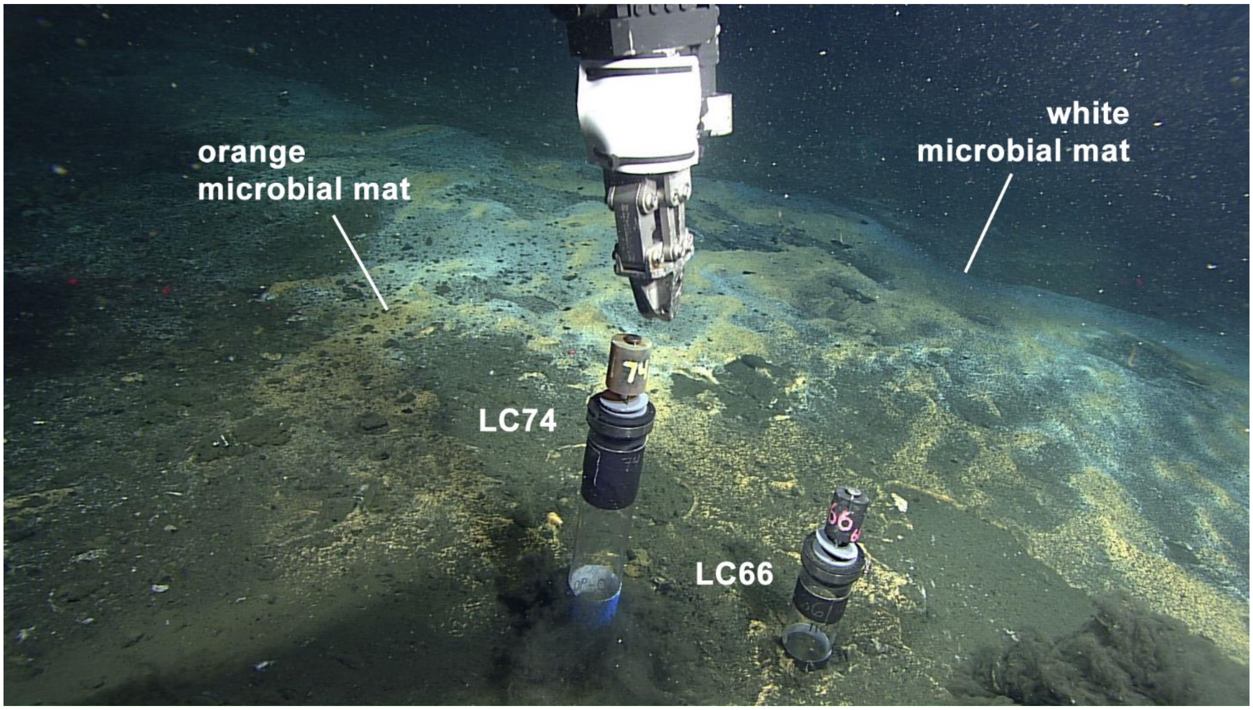


**Fig. S4** – *In situ* images of the sampling site at SMM 863 during the WF05-21 cruise. LC66 and LC74 were sampled over the same orange microbial mat. Core diameters shown are ~7 centimeters.

*Sediment Push Core Descriptions*

Sediment cores from all four sites in this study featured three primary material components: black to olive-colored mud; white bivalve shell fragments ranging from large, >cm-scale shards to mm-scale 'shell hash'; and hardened, grey-brown calcitic material ranging in size from <cm rubble to >cm nodules (Fig. S5). We also observed a mm-sized layer of microbial mat material overlying all the cores. Calcitic rubble was found at distinct depth intervals in the sediment at all four sites and was not exclusively associated with the bivalve shell fragments. Additionally, sites with long push core (LC, ~122 cm core barrel length) sampling (SMM 800-I and SMM 863) featured calcitic rubble and nodules as deep as 57 cmbsf and 48 cmbsf, respectively. The deeper cores did not feature nodules (>cm concretions) until 24-27 cmbsf, compared to the shallower nodules (3-12 cm cmbsf) recovered within the shorter push cores (PCs) at SMM 800-II and SMM 800-III. While nodules were observed in cores PC64 and PC43 (Fig. S5), these were not recovered for further analysis. Table S1 summarizes the site-specific, nodule-bearing sediment horizons included in this study.

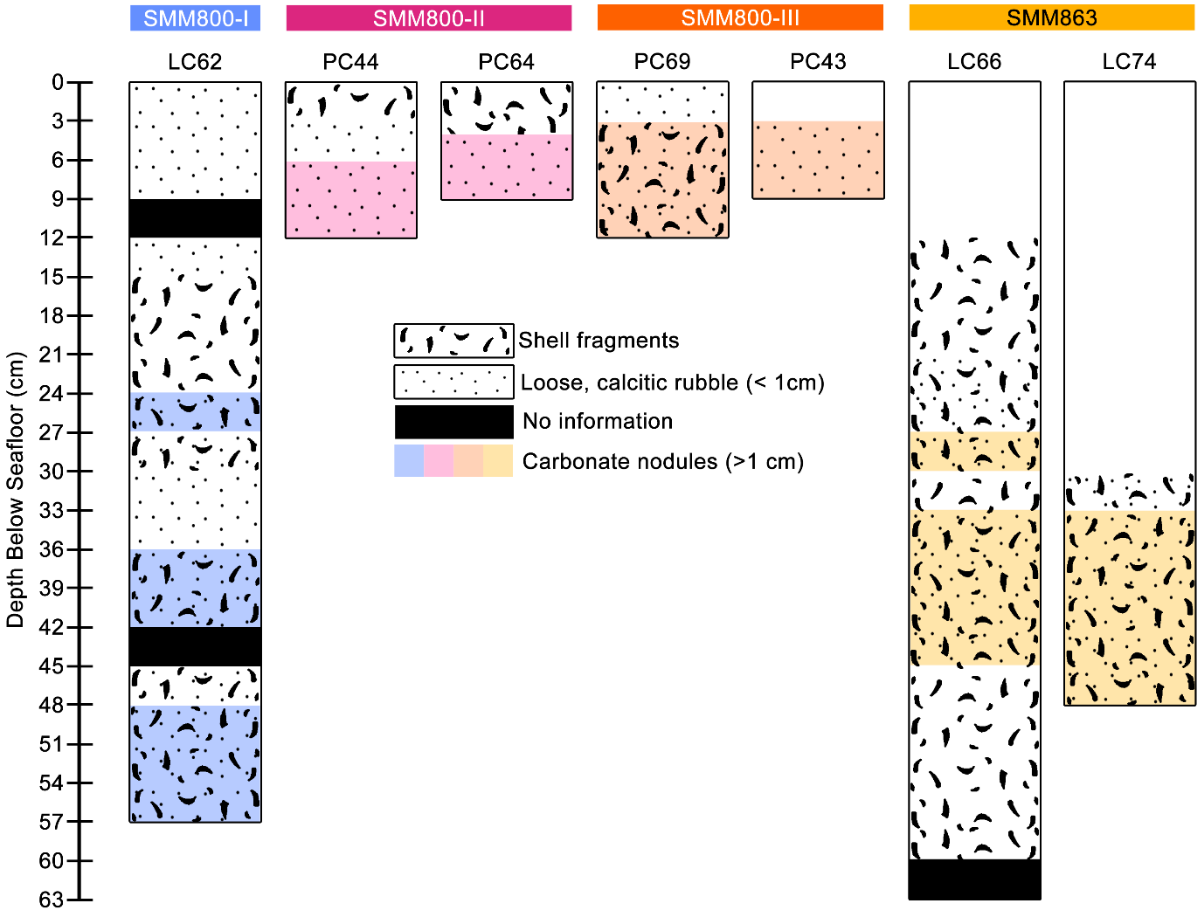


**Fig. S5** – Pseudo-stratigraphical characterizations of all the sediment push cores collected for this study, including shorter push cores (PC) and longer cores (LC). Here, we distinguish calcitic rubble from nodules on a size-basis, where the presence of concretions >1cm were considered nodules and anything smaller was excluded.

### Table S1

Nodule 'horizons' recovered for this study

| **Site** | **SMM 800 - Site I** | **SMM 800 - Site II** | **SMM 800- Site III** | **SMM 863** | |
|---|---|---|---|---|---|
| **Core ID** | LC62 | PC44 | PC69 | LC66 | LC74 |
| **Nodule Horizon (cmbsf)** | 24-27<br>36-42<br>48-51<br>51-57 | 6-9<br>9-12 | 3-12 | 36-39<br>39-42<br>42-45 | 33-39<br>42-45 |

### Geochemistry Supplemental Data

**Table S2**
Individual $\delta^{13}C$ for recovered nodules

| Site | Core ID | Depth (cmbsf) | $\delta^{13}C$ (‰) |
|---|---|---|---|
| SMM 800-I | LC62 | 24-27 | -30.49<br>-29.70 |
| | | 48-51 | -45.80<br>-46.25 |
| SMM 800-II | PC44 | 9-12 | -51.92<br>-51.84 |
| SMM 800-III | PC69 | 3-12 | -53.67<br>-53.92 |
| SMM 863 | LC74 | 33-39 | -31.85<br>-34.53 |
| | LC66 | 39-42 | -49.94<br>-49.15 |

*Note: values listed for each nodule horizon are replicate analyses from a single, representative nodule*

### 16S Sequencing Methods & Supplemental Data

*DNA Extraction, SSU rRNA Gene Amplification, and Illumina Sequencing*

During core sectioning on the WF05-21 and the WF02-20 cruises, 2mL of sediment from each core horizon were sampled and immediately stored at -80°C for later sequencing. During nodule extraction, rinsed nodules designated for sequencing were additionally treated to minimize sedimentary contamination according to the methods optimized in (4). Briefly, each nodule was rinsed in fresh, 0.22-µm-filtered, 1X PBS and sonicated at 8W for 45 seconds in fresh, 0.22-µm-filtered, 1X PBS. Rinsing and sonication was done three times using fresh 1X PBS each time before nodules were centrifuged in fresh, 0.22-µm-filtered, 1X PBS at 4,000g for 5 minutes and removed from the supernatant. After the 'decontamination' treatment, the nodules were finely ground with a sterile agate mortar and pestle. The mortar and pestle were washed with bleach, DI $H_2O$, and flame sterilized with ethanol prior to each use. DNA was extracted from 0.15-0.3g (wet weight) of thawed sediment or ground nodules with the DNeasy PowerSoil Pro Kit (Qiagen). The extraction protocol was modified for nodule samples to increase DNA yields according to methods developed by (5). After confirming satisfactory yields from sediment and nodules with a Qubit 2.0 fluorometer (Invitrogen), we performed a series of PCR amplifications for a 2-step Illumina sequencing strategy, detailed in (6). For this, we first performed a duplicate initial PCR amplification of the 16S rRNA gene using the 515f and 926r primer sets with Illumina adapters (515f; 5'-TCGTCGGCAGCGTCAGATGTGTATAAGAGACAG-GTGYCAGCMGCCGCGGTAA-3'; 926r; 5'-GTCTCGTGGGCTCGGAGATGTGTATAAGAGACAG-CCGYCAATTYMTTTRAGTTT-3', Illumina adapter show on the 5' end). The 515f and 926r primer set covers the V4 and V5 regions of the 16S rRNA gene and amplifies both bacterial and archaeal DNA, while excluding eukaryotic DNA. PCR reaction mixes were prepared with Q5 Hot Start High-Fidelity 2x Master Mix (New England Biolabs) in a 15 µL reaction volume according to manufacturer's directions with annealing conditions of 54°C for 30-35 cycles. With each PCR run, we ran a negative control using 1 µL of solution from a blank sample extracted with the PowerSoil Pro Kit instead of the DNA template and another negative control using 1 µL of DI $H_2O$. Following validation of the desired amplified sequence, duplicate samples were pooled and barcoded with Illumina Nextera XT index 2 primers that included unique 8-bp barcodes (P5; 5'-AATGATACGGCGACCACCGAGATCTACAC-XXXXXXXX-TCGTCGGCAGCGTC-3' and P7; 5'-CAAGCAGAAGACGGCATACGAGAT-XXXXXXXX-GTCTCGTGGGCTCGG-3'). Amplification with barcoded primers was done with Q5 Hot Start PCR mixture and 3 µL of the pooled product in a 30 µL reaction volume, annealed at 66°C, and cycled 11 times. After further validation, the barcoded PCR products were then combined in equimolar amounts into a single tube and 300uL of this pooled sample was run on a 1.5% low melt agarose gel (Fisher#BP165-25) and purified using Promega's Wizard SV Gel and PCR Clean-up System, Promega#A9281. Finally, amplicons were sequenced (2 x 300 bp) using the MiSeq Reagent Kit v3 (600-cycle) #MS-102-3003 on Illumina's MiSeq platform with the addition of 15-20% PhiX at Laragen (Culver City, CA, USA).

*Read Processing and Taxonomic Assignment*

At Laragen, the de-multiplexed forward and reverse reads were passed through a barcode filter that removed any reads with >1 bp mismatch. Filtered reads were then assigned a quality (Q) score, and adapter, index, and sequencing primer sequences were removed prior to arrival at the laboratory. Following hand-off of the paired-end sequences, further processing was performed on the reads with a Python-based pipeline. First, samples were filtered for the correct read length before the reads were trimmed to remove degenerate primer sequences using 'Cutadapt' (v2.9), (7). We kept reads within 10% of the expected read length after primer removal (between 252 and 308bp in length). Trimmed reads were then input into QIIME2 (v2020.11), (8), where they were filtered and denoised with a DADA2 pipeline (v1.14.1), (9). Filtering was accomplished in DADA2 by manually specified truncation of reads in order to remove low quality base calls while retaining enough overlap between forward and reverse reads for merging (12 nucleotides minimum). Because reads were compiled from 5 separate sequencing runs, 5 different truncation value pairs were used to account for run variability. Truncation values thus ranged from 242-267 for the forward reads and 156-226 for the reverse reads. After filtering, additional reads were removed based on a maximum expected error limit of 2 and a quality score threshold of 2. Chimeric sequence removal was also accomplished with DADA2, identified by consensus across samples. Finally, the DADA2 pipeline merged the remaining paired-end sequences.

ASVs with assigned taxonomy were generated from the merged, non-chimeric sequences in QIIME2 using the Silva V138.1 SSURef database feature classifier (10). Further thinning of the assigned reads was performed to remove ASVs unique to the negative controls from the dual-amplification steps, as well as singleton ASVs introduced by removing samples. Additionally, we used the 'decontam' R package's prevalence-based identification to remove likely contaminants based on the dual-amplicon negative controls, using a prevalence threshold of 0.5 (11). We also removed unassigned and Eukaryotic taxonomic assignments at the domain level, as well as any chloroplast taxonomic assignments to limit our focus to Bacterial and Archaeal diversity. After all processing was completed, the final data set had a mean sampling depth of 7148 reads. Samples with <1000 reads were dropped from the study.

*Phylum 16S rRNA gene Relative Abundances*

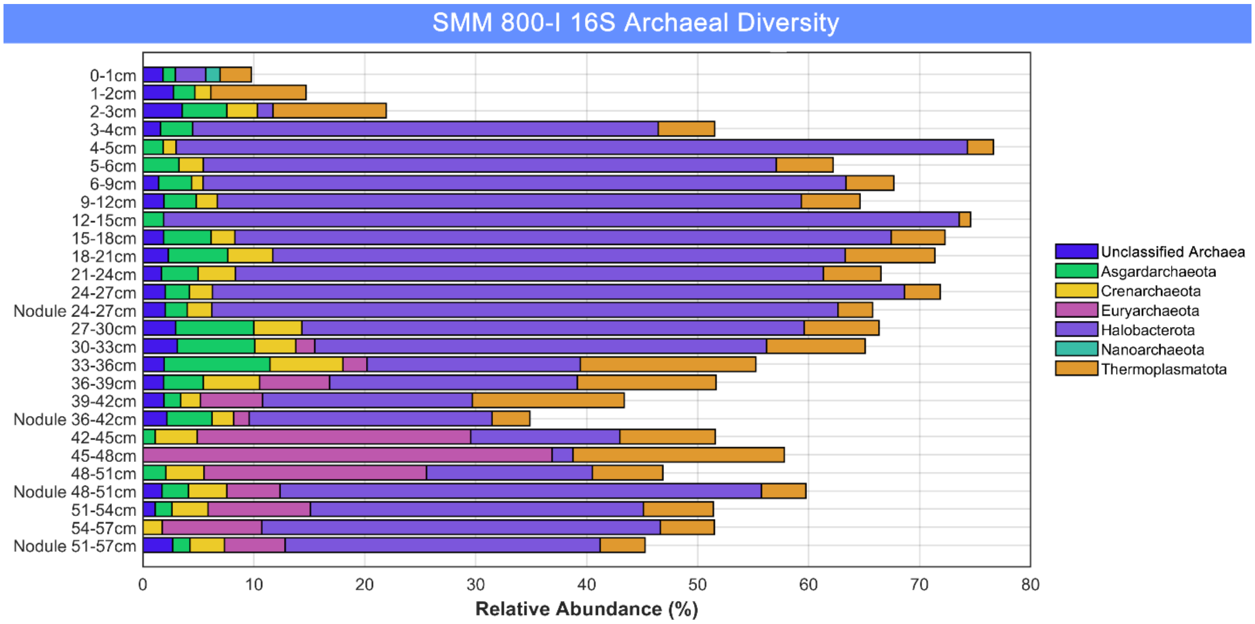


**Fig. S6** – Stacked bar charts summarizing the depth-resolved, phylum-level 16S archaeal abundances above 1% relative to total 16S abundances in LC62 at SMM 800-I.

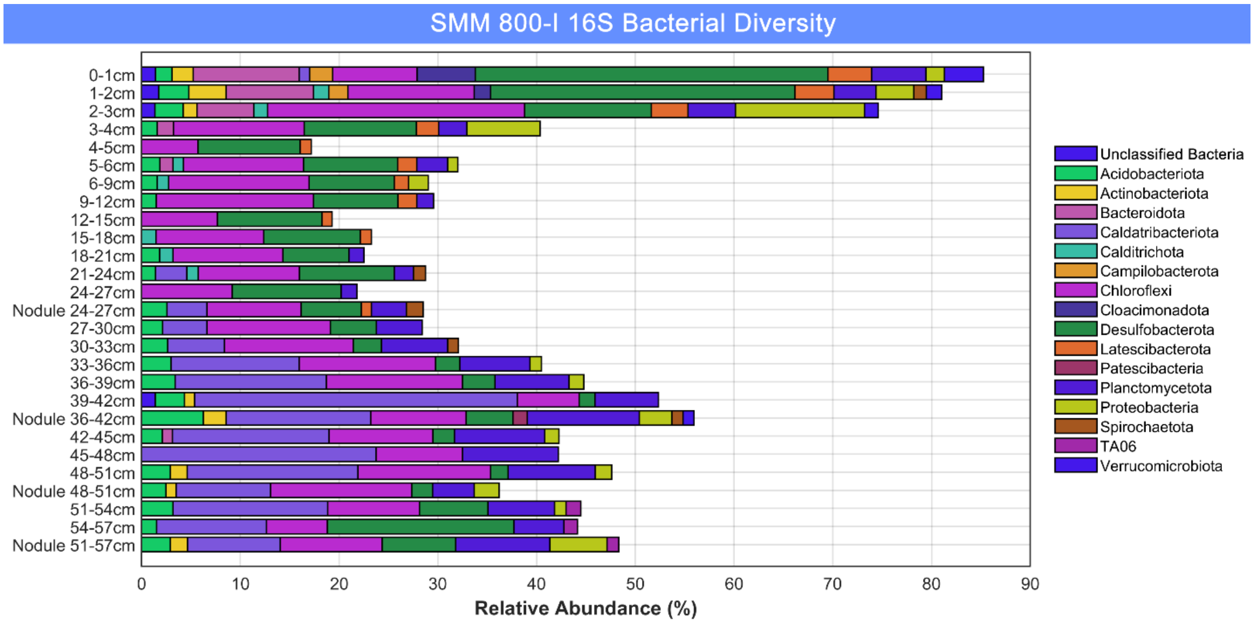


**Fig. S7** – Stacked bar charts summarizing the depth-resolved, phylum-level 16S bacterial abundances above 1% relative to total 16S abundances in LC62 at SMM 800-I.

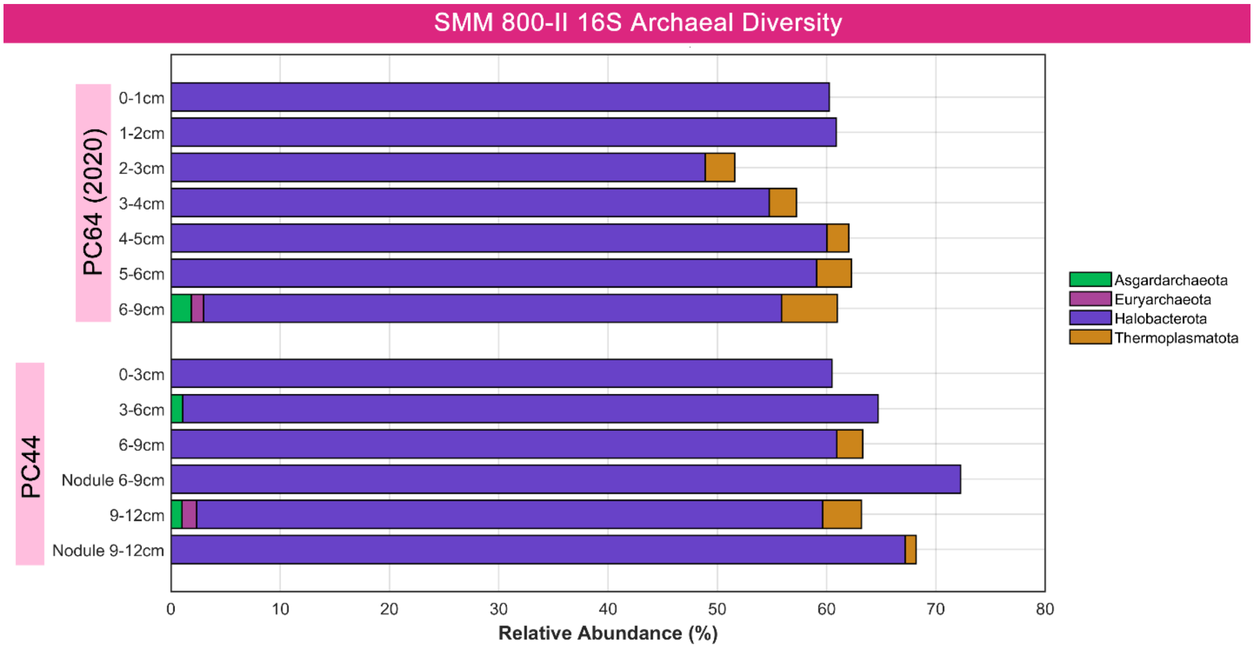


**Fig. S8** – Stacked bar charts summarizing the depth-resolved, phylum-level 16S archaeal abundances above 1% relative to total 16S abundances in PC64 and PC44 at SMM 800-II.

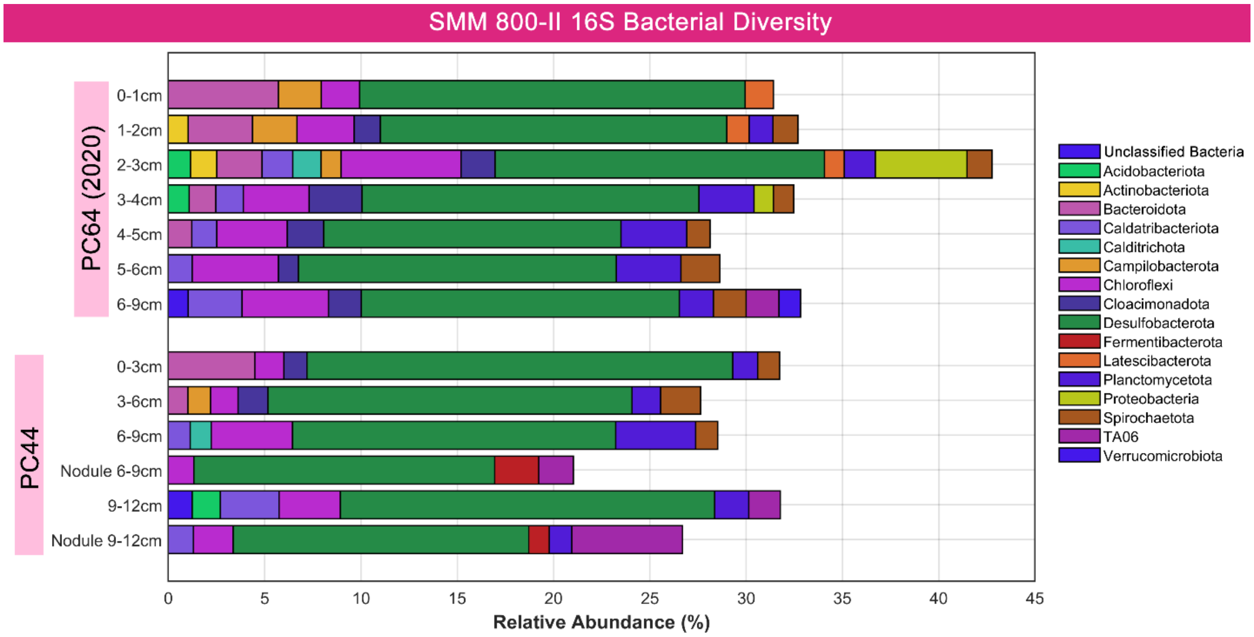


**Fig. S9** – Stacked bar charts summarizing the depth-resolved, phylum-level 16S bacterial abundances above 1% relative to total 16S abundances in PC64 and PC44 at SMM 800-II.

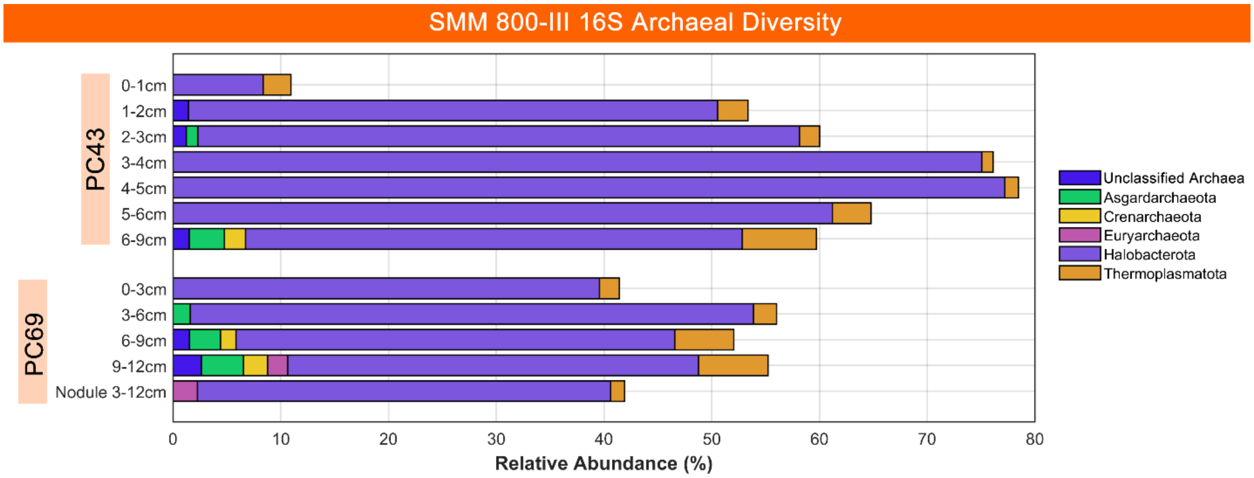


**Fig. S10** – Stacked bar charts summarizing the depth-resolved, phylum-level 16S archaeal abundances above 1% relative to total 16S abundances in PC43 and PC69 at SMM 800-III.

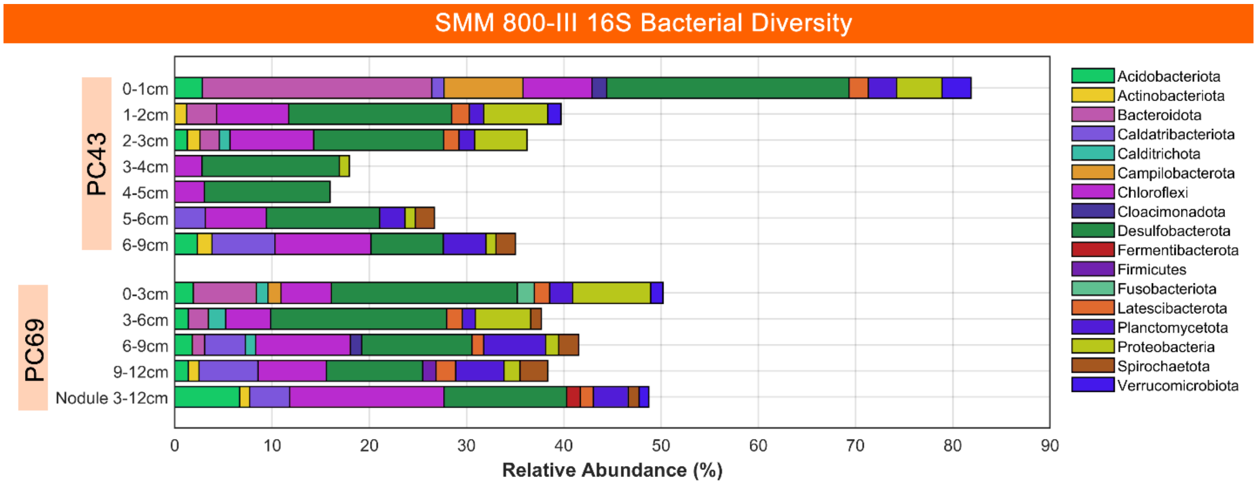


**Fig. S11** – Stacked bar charts summarizing the depth-resolved, phylum-level 16S bacterial abundances above 1% relative to total 16S abundances in PC43 and PC69 at SMM 800-III.

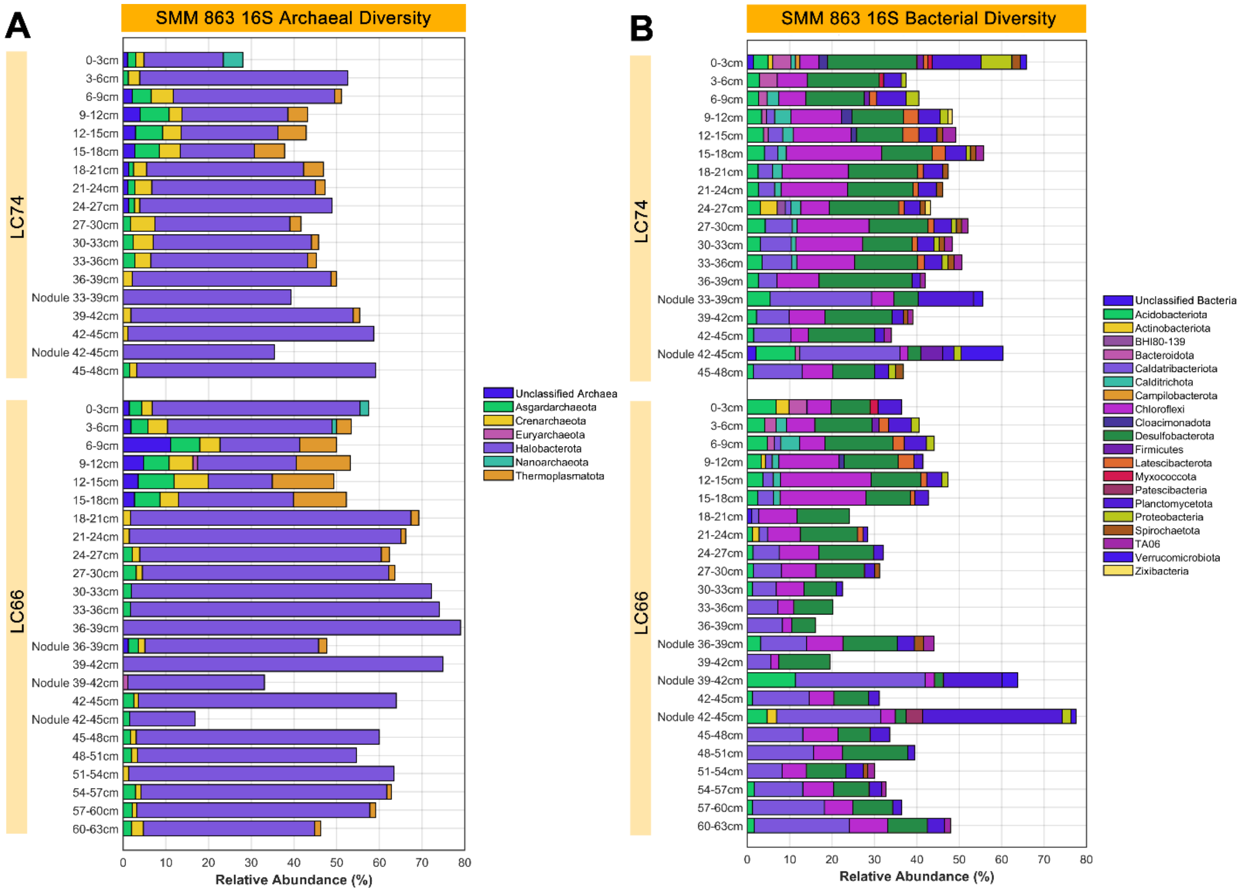


**Fig. S12** – Stacked bar charts summarizing the depth-resolved, phylum-level 16S A) archaeal and B) bacterial abundances above 1% relative to total 16S abundances in LC74 and LC66 at SMM 863.

*Diversity Index Calculations*

Both Shannon-Wiener and Inverse Simpson indices for archaeal and bacterial ASVs were calculated by sectioning the sequenced ASVs into bacterial and archaeal groups for each sample. The Shannon-Wiener indices were calculated per sample according to Eq. S1:

$$-\sum_{i=1}^{s} r_i \ln(r_i) \qquad (1)$$

where $r_i$ is the proportional relative abundance of each individual ASV ($i$) relative to the total number of archaeal or bacterial ASVs on a per sample basis ($s$). The Inverse Simpson indices were calculated per sample according to Eq. S2:

$$\frac{1}{\sum_{i=1}^{s} r_i^2} \quad (2)$$

where $r_i$ is the proportional relative abundance of each individual ASV ($i$) relative to the total archaeal or bacterial ASV pool on a per sample basis ($s$).

**Table S3**

Archaeal and bacterial diversity indices for recovered nodules and their host sediment horizons at each of the four sites in this study, calculated from domain-specific, 16S ASV proportional abundances

<table>
<tr><th rowspan="3">Site</th><th rowspan="3">Core ID</th><th rowspan="3">Core Horizon (cmbsf)</th><th colspan="4">Shannon-Wiener Index</th><th colspan="4">Inverse Simpson Index</th></tr>
<tr><th colspan="2">Archaea</th><th colspan="2">Bacteria</th><th colspan="2">Archaea</th><th colspan="2">Bacteria</th></tr>
<tr><th>Sediment</th><th>Nodule</th><th>Sediment</th><th>Nodule</th><th>Sediment</th><th>Nodule</th><th>Sediment</th><th>Nodule</th></tr>
<tr><td rowspan="6">SMM 800-I</td><td rowspan="6">LC62</td><td>24-27</td><td>2.3</td><td>2.9</td><td>3.8</td><td>4.7</td><td>4.0</td><td>6.1</td><td>29.9</td><td>64.7</td></tr>
<tr><td>36-39</td><td>3.1</td><td rowspan="2">2.9</td><td>3.9</td><td rowspan="2">5.1</td><td>13.8</td><td rowspan="2">5.3</td><td>20.6</td><td rowspan="2">48.3</td></tr>
<tr><td>39-42</td><td>2.7</td><td>3.0</td><td>10.7</td><td>8.0</td></tr>
<tr><td>48-51</td><td>2.6</td><td>2.7</td><td>4.0</td><td>4.2</td><td>5.9</td><td>5.0</td><td>22.3</td><td>29.2</td></tr>
<tr><td>51-54</td><td>3.4</td><td rowspan="2">3.4</td><td>4.1</td><td rowspan="2">4.7</td><td>17.2</td><td rowspan="2">15.6</td><td>19.8</td><td rowspan="2">38.8</td></tr>
<tr><td>54-57</td><td>3.1</td><td>3.3</td><td>12.3</td><td>13.8</td></tr>
<tr><td rowspan="2">SMM 800-II</td><td rowspan="2">PC44</td><td>6-9</td><td>2.3</td><td>1.7</td><td>4.1</td><td>3.5</td><td>5.2</td><td>2.7</td><td>32.9</td><td>12.1</td></tr>
<tr><td>9-12</td><td>2.6</td><td>1.8</td><td>3.6</td><td>3.3</td><td>7.5</td><td>3.0</td><td>14.9</td><td>11.0</td></tr>
<tr><td rowspan="3">SMM800-III</td><td rowspan="3">PC69</td><td>3-6</td><td>2.6</td><td rowspan="3">2.3</td><td>4.5</td><td rowspan="3">4.4</td><td>6.9</td><td rowspan="3">5.6</td><td>48.3</td><td rowspan="3">25.1</td></tr>
<tr><td>6-9</td><td>3.3</td><td>4.8</td><td>11.2</td><td>61.9</td></tr>
<tr><td>9-12</td><td>3.1</td><td>4.3</td><td>13.9</td><td>43.5</td></tr>
<tr><td rowspan="6">SMM 863</td><td rowspan="3">LC66</td><td>36-39</td><td>1.6</td><td>2.7</td><td>3.4</td><td>4.7</td><td>2.1</td><td>5.0</td><td>15.0</td><td>43.8</td></tr>
<tr><td>39-42</td><td>1.9</td><td>2.1</td><td>3.3</td><td>2.9</td><td>2.4</td><td>3.4</td><td>13.4</td><td>8.9</td></tr>
<tr><td>42-45</td><td>2.5</td><td>2.2</td><td>3.9</td><td>3.4</td><td>3.8</td><td>3.4</td><td>22.6</td><td>9.3</td></tr>
<tr><td rowspan="3">LC74</td><td>33-36</td><td>2.2</td><td rowspan="2">2.1</td><td>4.3</td><td rowspan="2">3.6</td><td>4.3</td><td rowspan="2">2.8</td><td>41.8</td><td rowspan="2">13.7</td></tr>
<tr><td>36-39</td><td>2.7</td><td>4.2</td><td>4.7</td><td>18.2</td></tr>
<tr><td>42-45</td><td>2.2</td><td>1.4</td><td>4.0</td><td>3.5</td><td>3.1</td><td>1.8</td><td>21.7</td><td>9.7</td></tr>
</table>

Neither index demonstrated a correlation with the number of bacterial or archaeal ASVs recovered.

*Community Statistical Analysis*

Non-metric multi-dimensional scaling (NMDS) ordination was done in MATLAB (v.2022a) using the open-source Fathom toolbox designed by (12) at USF's College of Marine Science. The Bray-Cutris dissimilarity matrix was first calculated using all ASV total relative abundances (on a per sample basis) with Fathom's `f_braycurtis` function. The NMDS was generated using MATLAB's `mdscale` with the 'stress' criterion specified, which normalizes stress by the sum of squares of the interpoint distances.

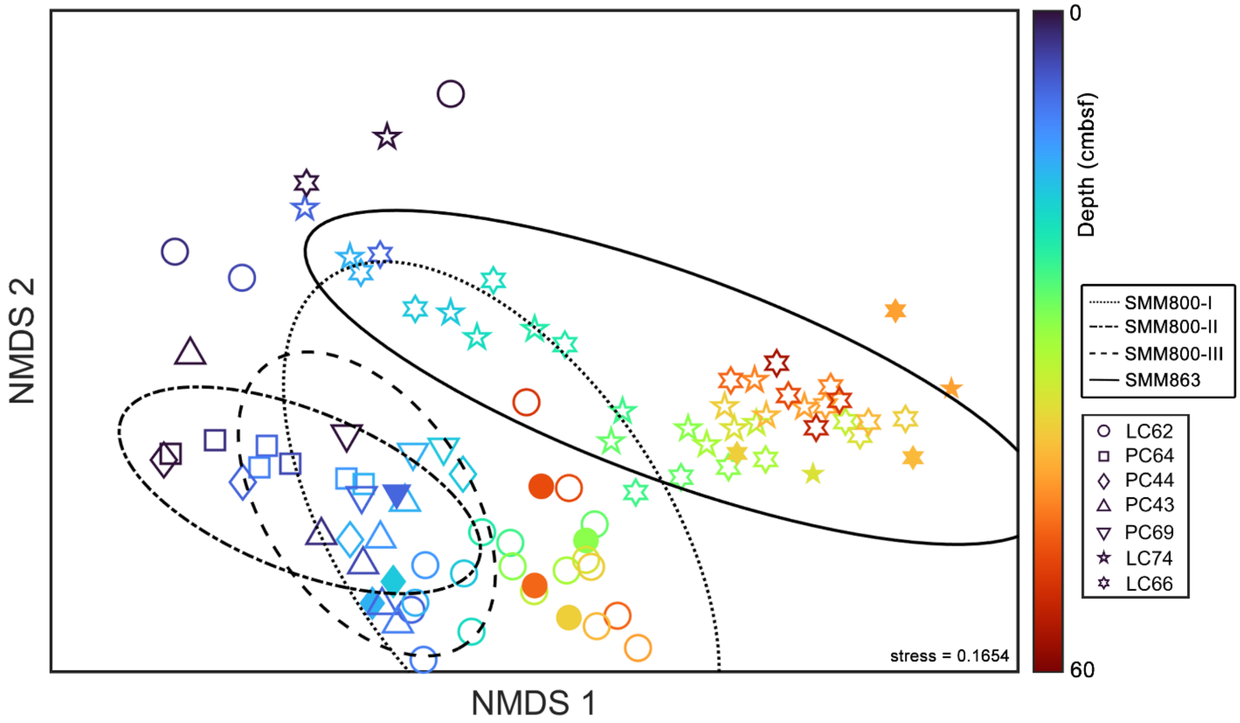


**Fig. S13 –** Nonmetric multidimensional scaling (NMDS) ordination of all 16S ASV relative abundances in the sediment horizons and their nodules, where collected. Empty and filled in markers correspond to horizon-specific sediment and nodule communities, respectively. The shape colors denote depth in cm below seafloor (cmbsf). The four sampling sites (SMM 800-I, SMM 800-II, SMM 800-III, and SMM 863) are indicated with 95% confidence interval ellipses.

### **Fluorescence *In Situ* Hybridization**

**Table S4**
FISH hybridization probe mix 1 (40% formamide, per 101-µL)

| Reagent | Amount (in µL) |
|---|---|
| 5M NaCl | 18 |
| 1M Tris-HCl (pH 8) | 2 |
| EUB338 + EUB338 II + EUB338 III (each at 50 ng/µL in PCR-quality, ultrapure DI $H_2O$) | 10 |
| ANME1-350 (50 ng/µL in PCR-quality, ultrapure DI $H_2O$) | 10 |
| ANME1-728 (50 ng/µL in PCR-quality, ultrapure DI $H_2O$) | 10 |
| ARCH-915 (50 ng/µL in PCR-quality, ultrapure DI $H_2O$) | 10 |
| Formamide | 40 |
| 1% SDS (by weight in sterile, ultrapure DI $H_2O$) | 1 |

**Table S5**
FISH hybridization wash (for 40% formamide hybridization, per 2-mL)

| Reagent | Amount (in µL) |
|---|---|
| 5M NaCl | 18.4 |
| 1M Tris-HCl (pH 8) | 40 |
| 0.5M EDTA | 20 |
| Sterile, ultrapure DI $H_2O$ | 1921.6 |

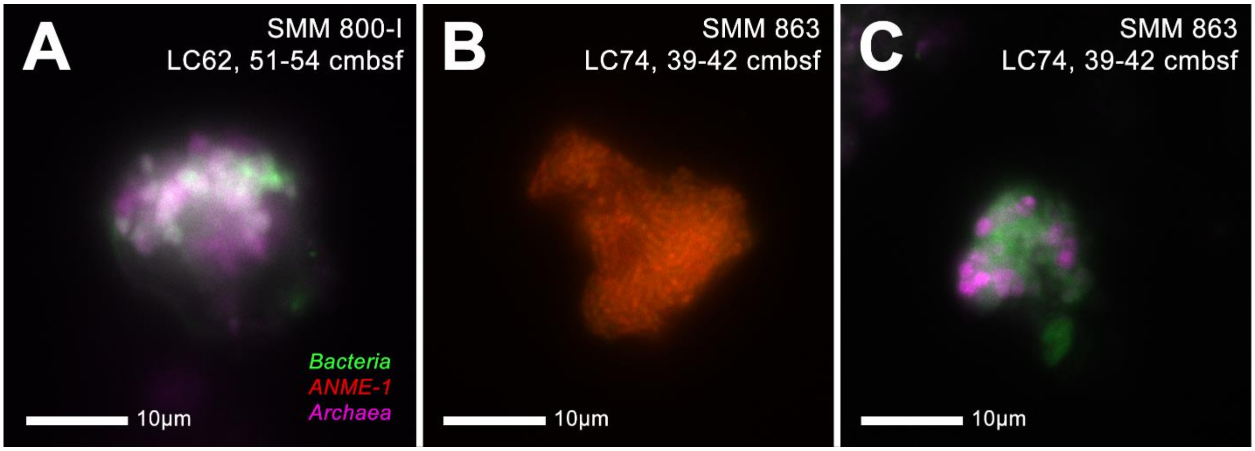


**Fig. S14** – Observations of aggregate morphologies extracted from deeper sediment horizons at SMM 800-I and SMM 863, including those consistent with ANME-SRB consortia and positive fluorescence in-situ hybridization (FISH) signal from archaeal (ARCH915, in pink; ANME1-350, ANME-728, in red) and bacterial (EUB338, EUB338 II, EUB338 III, in green) probes.

### Single Cell Translational Activity Measurements with BONCAT

Recovered nodules were incubated in artificial seawater (ASW) medium. The ASW was prepared according to the amounts listed below, modified from (13) and (14) using Wolin' vitamin solution from DSMZ 141 medium. Notable changes from other ASW media include reducing the as the buffer and modified salt concentrations to ensure $\Omega_{calcite}$ and $\Omega_{aragonite}$ were close to 1 (minimizing the impact of carbonate precipitation/dissolution, see (13)). Reagents were dissolved in deionized water at room temperature, then 0.2-μm filter-sterilized and sparged with $N_2$ until anoxic. Media was chilled at 4°C overnight prior to use.

**Table S6**

Artificial seawater media (per 1-L DI $H_2O$)

| Reagent | Amount | Final Concentration [mM] |
|---|---|---|
| $MgCl_2 \cdot 6H_2O$ | 9.47 g | 46.60 |
| $CaCl_2 \cdot 2H_2O$ | 1.47 g | 10.00 |
| NaCl | 25.70 g | 439.77 |
| KCl | 0.52 g | 7.00 |
| $Na_2SO_4$ | 1.42 g | 10.00 |
| $K_2HPO_4$ | 0.17 g | 1.00 |
| $NH_4Cl$ | 0.11 g | 2.00 |
| HEPES Buffer (250mM, pH 7.5) | 100.0 mL | 25.00 |
| Vitamins DSMZ 141 (1000x) | 1.0 mL | *varies per vitamin* |
| $NaHCO_3$ (1M) | 5.0 mL | 5.00 |
| *Se/W Solution as in (14), containing per 1L:* | | |
| $Na_2SeO_3$ | 0.017 g | - |
| $Na_2WO_4 \cdot 2H_2O$ | 0.033 g | - |
| *Trace Element Solution as in (14), containing per 1L:* | | |
| Nitrilotriacetic acid (NTA) | 0.15 g | - |
| $MnCl_2 \cdot 4H2O$ | 0.61 g | - |
| $CoCl_2 \cdot 6H2O$ | 0.42 g | - |
| $ZnCl_2$ | 0.09 g | - |
| $CuCl_2 \cdot 2H_2O$ | 0.007 g | - |
| $AlCl_3$ | 0.006 g | - |
| $H_3BO_3$ | 0.01 g | - |
| $Na_2MoO_4 \cdot 2H_2O$ | 0.03 g | - |
| $SrCl_2 \cdot 6H_2O$ | 0.01 g | - |
| NaBr | 0.01 g | - |
| KI | 0.07 g | - |
| $FeCl_3 \cdot 6H_2O$ | 0.5 g | - |
| $NiCl_2 \cdot 6H_2O$ | 0.025 g | - |

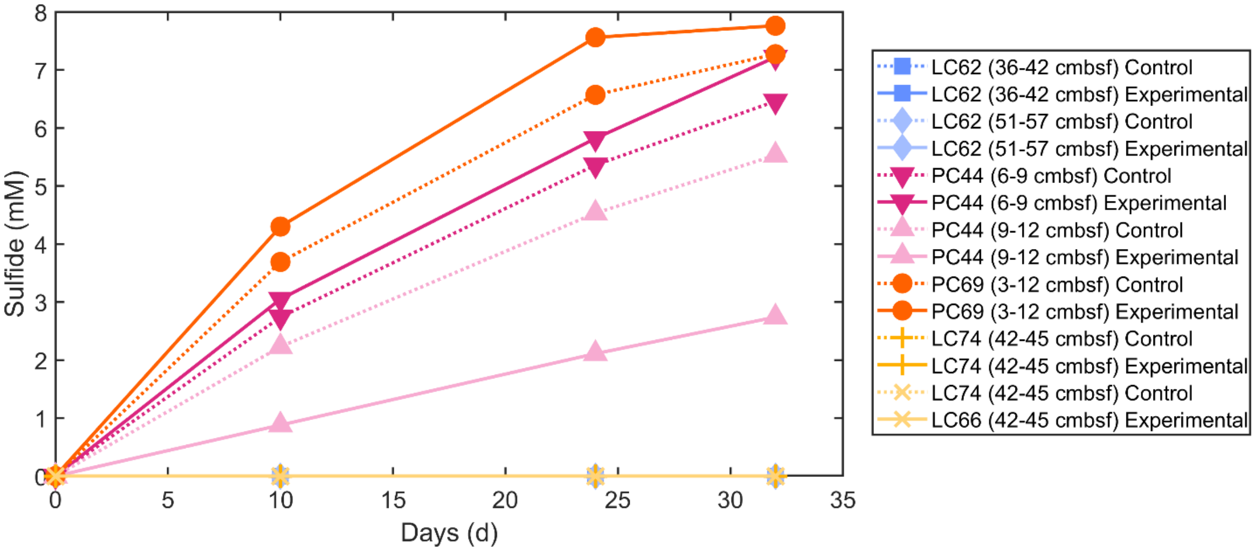


**Fig. S15** – Sulfide production in the 5-week period after separation of incubated nodules into control and experimental groups, but before addition of HPG, as observed by dissolved sulfide in media and measured by Cline assay. Comparable sulfide production attests to sulfate-coupled AOM activity in both bottles. Notably, both control and experimental groups for nodules from SMM 800-I and SMM 863 did not produce a detectable amount of sulfide.

**Table S7**

BONCAT azide dye solution (per 250-µL)

| Reagent | Amount (in µL) |
|---|---|
| 100mM Aminoguanidine (in ultrapure DI $H_2O$) | 12.5 |
| 100mM Ascorbate (in 1X PBS) | 12.5 |
| 20mM $CuSO_4$ (in ultrapure DI $H_2O$) | 1.25 |
| 50mM BTTAA (in ultrapure DI $H_2O$) | 2.5 |
| 10mM Oregon Green 488 Azide dye (VectorLabs) | 0.5 |
| 1X PBS | 221 |

Note: A dye ‘pre-mix’ containing the azide dye, BTTAA, and $CuSO_4$ was incubated under Ar in the dark for 3 minutes prior to the addition of the rest of the reagents).

**Table S8**

FISH hybridization probe mix 2 (35% formamide, per 100-µL)

| Reagent | Amount (in µL) |
|---|---|
| 5M NaCl | 18 |
| 1M Tris-HCl (pH 8) | 2 |
| EUB338 + EUB 338 II + EUB338 III | 10 |
| (each at 50 ng/µL in PCR-quality, ultrapure DI $H_2O$) | |
| ARCH-915 (50 ng/µL in PCR-quality, ultrapure DI $H_2O$) | 10 |
| Formamide | 35 |
| 1% SDS (by weight in sterile, ultrapure DI $H_2O$) | 1 |
| PCR-quality, ultrapure DI $H_2O$ | 24 |
| 5M NaCl | 18 |

**Table S9**
FISH hybridization wash (for 35% formamide hybridization, per 2-mL)

| Reagent | Amount (in µL) |
|---|---|
| 5M NaCl | 28 |
| 1M Tris-HCl (pH 8) | 40 |
| 0.5M EDTA | 20 |
| Sterile, ultrapure DI $H_2O$ | 1912 |

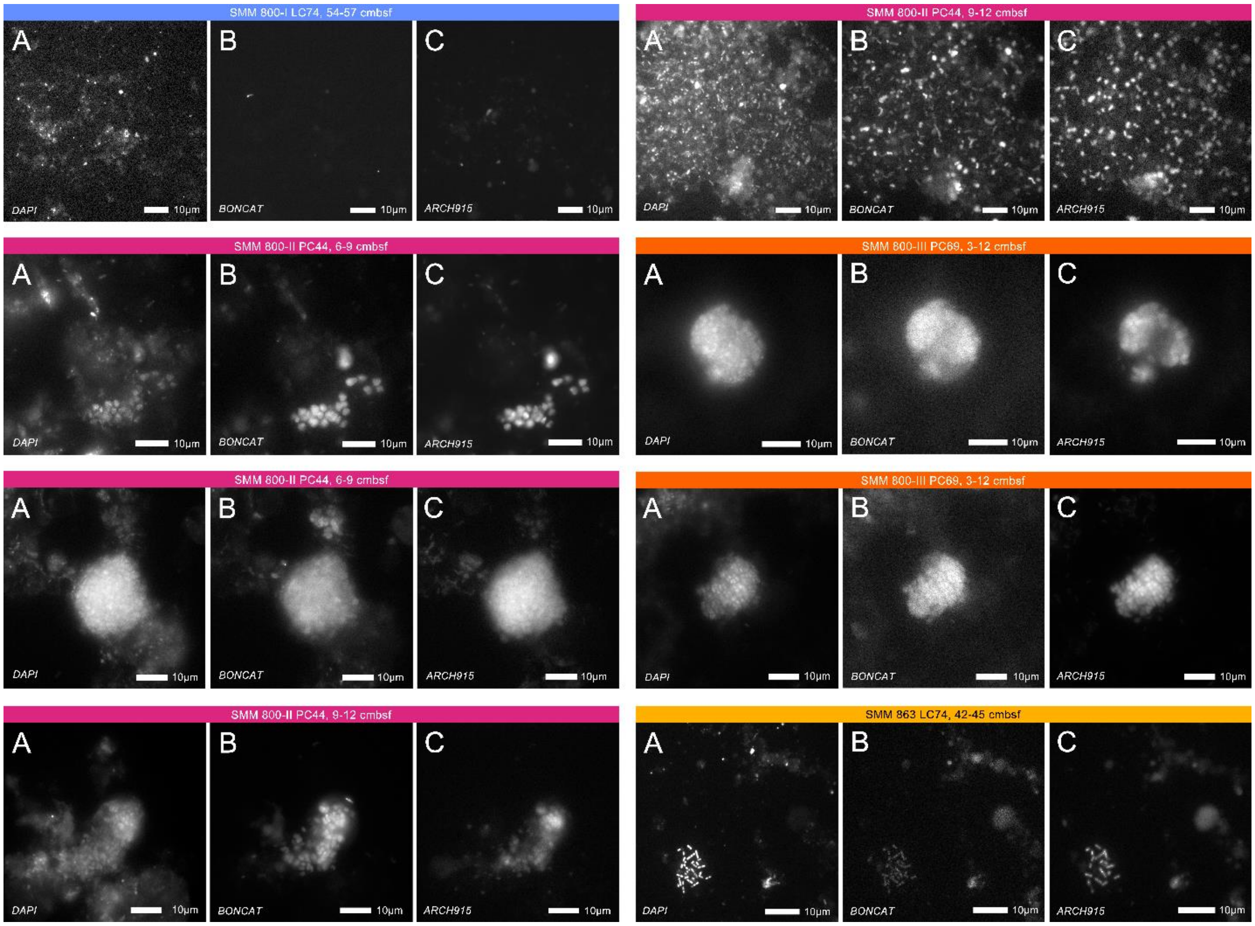


**Fig. S16** – Individual fluorescent channels from BONCAT-FISH observations of cells from incubated nodules shown in Fig. 13 where A) corresponds to DAPI signal; B) is signal from BONCAT click dye Oregon Green; C) reflects signal from archaeal probe ARCH915.
